# Polypharmacological Re-programming of Tumor-associated Macrophages Restores Antitumor Immunity

**DOI:** 10.1101/2021.04.10.439265

**Authors:** Nao Nishida-Aoki, Taranjit S. Gujral

**Affiliations:** Human Biology Division, Fred Hutchinson Cancer Research Center, Seattle, WA, USA; Department of Pharmacology, University of Washington, Seattle, WA, USA

**Keywords:** Tumor-associated macrophage, Breast cancer, Tumor microenvironment, Kinase inhibitor screening, Cell-cell interaction, Actin filament organization, BMS-794833, Conditioned medium

## Abstract

Tumor-associated macrophages (TAMs) are an important component of the tumor microenvironment (TME), and they promote tumor progression, metastasis, and resistance to therapies. However, although TAMs represent a promising target for therapeutic intervention, the complexity of TME has made the study of TAMs challenging. Here, we established a physiologically-relevant *in vitro* TAM polarization system that recapitulates TAM pro-tumoral activities. We used this system for phenotypic kinase inhibitor screening and identified a multi-targeted compound BMS-794833 as the most potent inhibitor of TAM polarization. BMS-794833 decreased pro-tumoral properties of TAMs and suppressed tumor growth in mouse triple-negative breast cancer models. The effect of BMS-794833 was not dependent on its primary targets (MET and VEGFR2) but on its effect on multiple signaling pathways, including focal adhesion kinases, SRC family kinases, STAT3, and p38 MAP kinases. Our study underlines the efficacy of polypharmacological strategies in re-programming complex signaling cascades activated during TAM polarization.

## Introduction

Solid tumors comprise heterogeneous populations of cancer and non-cancerous cells that interact through direct contact and secreted factors, thus establishing tumor microenvironment (TME). TME alters the behavior of non-cancerous cells, resulting in phenotypes with tumor-supportive properties. For example, tumor cells and immune cells that reside within TME release a number of anti-inflammatory and pro-angiogenic factors that result in pro-tumoral polarization of tumor-associated macrophages (TAMs), one of the abundant types of immune cells found in TME. Thus, polarized TAMs can further promote tumor progression by stimulating proliferation, invasion, angiogenesis, immunosuppression, metastasis, endowing resistance against chemo- and radiotherapy, and decreased efficacy of immunotherapy (Cassetta and Pollard, 2018; Xiang et al., 2021). Further, clinical evidence supports that patients with higher TAM infiltration result in poor prognosis in several cancers(Jung et al., 2015). Given their role in tumor progression and the clinical evidence, TAMs have been proposed as a promising therapeutic target (Bingle et al., 2002; Fridman et al., 2017; Zhao et al., 2017). However, physiological TAMs are scarce and difficult to study using existing experimental strategies. Therefore, mechanisms that drive TAM pro-tumor polarization, as well as potential targeting opportunities for drug development remain incompletely understood.

Accumulated evidence suggests that TAMs are highly plastic and dynamic cells with complex signaling networks optimized to rapidly adjust the phenotype in response to external stimuli(Irey et al., 2019). Therefore, TAMs exhibit heterogenous(Azizi et al., 2018) and continuum phenotypes between inflammatory and anti-inflammatory(Movahedi et al., 2010). One strategy to model TME *in vitro* is to employ tumor conditioned medium (CM) of cancer cells, which has been shown to contain a near-complete set of secreted factors and metabolites found in TME and has been used in several TAM studies aimed at improving our understanding of TAM biology (Benner et al., 2019; Cabanel et al., 2015; Chen et al., 2017; Solinas et al., 2010; Su et al., 2014). However, quantitative, phenotypic drug screening assays aimed at TAMs have not been described before, thus limiting the ability to discover compounds with potential therapeutic benefits in this context. To address this need, we established an *in vitro* quantitative TAM polarization model using human monocyte cell line THP-1 and CM from multiple cancer models. Using cellular elongation measurement as a quantitative measure of TAM polarization, we screened a library of 85 kinase inhibitors targeting most of the human kinome. We identified BMS-7948933, a dual-targeting inhibitor of c-MET and VEGFR2, as the most potent blocker of cellular elongation and pro-tumoral function of TAMs, and demonstrated therapeutic effects on mouse triple-negative breast tumor models. Surprisingly, we found the TAM inhibitory function of BMS-794833 does not involve c-MET or VEGFR2 but a range of other targets that include focal adhesion kinases (FAK) and cytoskeletal-related proteins, SRC family kinases, STAT3, and p38 MAP kinases. We also observed that targeting a single pathway exhibited little to no effect on TAM polarization, whereas a combination of STAT3 inhibitor and LIMK (actin polymerization regulator) inhibitor exhibited an additive effect on TAM polarization. Our study underlines the complex regulation of signaling pathways during TAM polarization, and the necessity of concomitant blocking of multiple signaling pathways to re-program pro-tumoral TAM function.

## Material and Methods

### Cell lines and culture

Cell lines including 4T1, CT26, Py8119, THP-1, and Jurkat cells were purchased from American Type Culture Collection (ATCC). 4T1 and CT26 cells were maintained in RPMI1640 media (Gibco by Thermo Fisher Scientific, Waltham, MA, USA) supplemented with 10% (v/v) fetal bovine serum (FBS), 1% Penicillin-Streptomycin (P/S, final concentration: 100 units/ml of penicillin, 100 µg/ml of streptomycin) (Gibco). 4T1 cells with nuclear-localized GFP (4T1-nucGFP) were established by introducing NucLight Green Lentivirus Reagent (Essen BioScience by Sartorius, Goettingen, Germany) under the selection of 2 µg/ml puromycin. Py8119 cells were maintained in F-12K medium (ATCC) with 5% FBS. THP-1 cells were cultured in RPMI1640 media supplemented with 10% FBS, 1% P/S, 1 mM sodium pyruvate (Lonza, Basel, Switzerland), and 55 nM β-mercaptoethanol (Gibco). Jurkat cells were maintained in RPMI1640 media with 10% FBS, 1% P/S, 10 mM HEPES (Santa Cruz Biotechnology, Dallas, TX, USA), and 1 mM sodium pyruvate. B16.F10.Ova mouse melanoma cells (kindly provided by Dr. Anthony Rongvaux, Fred Hutchinson Cancer Research Center (FHCRC)) and Panc02 mouse pancreatic ductal carcinoma cells (kindly provided by Dr. Partecke, University of Greifswald) were cultured in DMEM (Corning by MilliporeSigma, Burlington, MA, USA) supplemented with 10% FBS and 1% P/S. MC38 cells (Kerafast, Boston MA, USA), was maintained in DMEM with 10% FBS, 1% P/S, 2 mM L-glutamine (Gibco), 1× non-essential amino acids (Gibco), 1 mM sodium pyruvate, 10 mM HEPES, and 50 µg/ml gentamycin. HMLE and HMLE-Ras (kindly provided by Dr. Robert Weinberg, Whitehead Institute) were cultured with DMEM/F-12 (Gibco), 10% FBS, 1% P/S, 20 ng/ml mouse EGF (BioLegend, San Diego, CA, USA), 10 µg/ml bovine insulin (Sigma-Aldrich by MilliporeSigma), and 50 nM hydrocortisone (Sigma). All cells were cultured in humidified 37°C incubators with 5% CO_2_ atmosphere.

### Collection of conditioned medium (CM)

For the collection of CM from tumor tissue and cancer cells, Tumor slice culture (TSC) medium(Nishida-Aoki et al., 2020), composed of William’s medium E (Gibco), 12 mM Nicotinamide (Sigma-Aldrich), 150 nM ascorbic acid (Sigma-Aldrich), 2.25 mg/ml sodium bicarbonate (Corning), 20 mM HEPES, 50 mg/ml glucose (Sigma-Aldrich), 1 mM sodium pyruvate, 2 mM L-glutamine, 1% ITS (Fisher Scientific), 20 ng/ml EGF, 40 IU/ml penicillin and 40 µg/ml streptomycin, was used. Cancer cells grown to subconfluent were washed with PBS twice and incubated in TSC medium for 1-2 days. For the collection of CM from tumor tissues, tumors were cut into 4-5 mm cubes, washed with PBS twice, and incubated with 10 ml of TSC medium per 1 g tumor for one day. After collecting the spent medium, fresh TSC medium was added to the tumor pieces to collect the second pool. The collected medium was centrifuged at 2,000 ×g for 10 min, and the supernatant was used as CM. TSC medium without conditioning was used as an experimental control. For the collection of TAM CM, TAMs polarized for 3 days were washed with PBS twice, then incubated for 24 h in RPMI1640 medium for THP-1 maintenance. The supernatant was collected and centrifuged at 2,000 ×g for 10 min, to remove cell debris. RPMI1640 medium for THP-1 culture without conditioning was used for experimental control.

### TAM polarization model with cancer CM

THP-1 cells were differentiated into macrophages by inducing with phorbol 12-myristate 13-acetate (PMA, LC Laboratories, Woburn, MA, USA) at 25 ng/ml for one day. The cell densities used were; 1.3-1.5×10^4^/well for 96-well plate, 4×10^5^ cells/well for 6-well plate, 2.4×10^6^ cells for 10 cm dish. The medium was replaced with medium containing cancer CM at 25-50% (v/v) with volume adjustment to 50% with TSC medium, and 50% (v/v) RPMI1640 for THP-1 cell culture. Cellular elongation was measured with NeuroTrack analysis software accompanied with IncuCyte Zoom Live Imaging system (Sartorius).

### Kinase inhibitor screening

THP-1 cells were seeded at 1.5×10^4^ cells per well of 96-well plates with PMA at 25 ng/ml. After a day, the medium was replaced to CM mix (25% 4T1 cell CM, 25% TSC medium, with 50% RPMI1640 for THP-1 culture, YOYO-3 at 1:10,000 dilution) together with kinase inhibitors at 8 serial concentrations of 10, 3.3, 1.1, 0.37, 0.12, 0.04, 0.01, and 0 µM with triplicates. The 85 inhibitors tested are listed in Table S1. All small molecules were constituted in DMSO for the stock solution, and DMSO (0.1%, up to 1% based on the volume of inhibitor solution) was supplemented as vehicle control. A red fluorescent viability dye, YOYO-3 (Thermo Fisher Scientific) was supplemented to the culture to detect cellular death. The phase contrast and red fluorescent images were taken every 2 hours over 3-5 days using IncuCyte Zoom instrument. The target kinase profiles of BMS-794833 in acellular system were described in Rata, et al.(Rata et al., 2020).

### Coculture of TAM and cancer cells

THP-1 cells at 1.5×10^4^ cells per well of 96-well plate were induced with PMA at 25 ng/ml and polarized for 3 days with the indicated tumor CM with or without inhibitors. The TAMs were washed with PBS twice, and 4T-nucGFP cells were seeded at 3×10^3^ cells per well density with RPMI1640 supplemented with 0.5% FBS. The cell numbers of 4T1 were counted based on nuclear GFP using the software accompanied with IncuCyte Zoom.

### Tube formation assay

Tube formation assay of endothelial cells was performed using IncuCyte Angiogenesis 96-well PrimeKit Assay (Sartorius) with modifications. For the angiogenesis assay of TAM CM, the cells were seeded following the manufacturer’s protocol. Three days post seeding, TAM CM was supplemented at 50% concentration diluted in the assay medium included in the kit. The cells were incubated for 6 days with a medium replacement on day 2 and 5 during the culture. The plate was scanned and analyzed with IncuCyte Zoom. For assay of BMS-794833-treated TAM CM, normal human dermal fibroblast (NHDF) and GFP-expressing human umbilical vein endothelial cells (GFP-HUVECs) were minimally expanded. NHDF were seeded at 1×10^4^ cells per well of 96-well plate with DMEM supplemented with 10% FBS and 1% P/S and incubated for 2 days till confluent. GFP-HUVECs were seeded at 2×10^3^ cells per well in EGM-2 on top of the NHDF layer. On the next day, the medium was replaced with 50% TAM CM diluted in EGM-2, and cells were incubated for 6 days. A half volume of medium was replated every 2 days. The plate was scanned and analyzed with IncuCyte S3.

### Jurkat migration assay

Jurkat cells were washed with PBS and seeded onto a 24-well transwell culture insert with 3 µm pores (Celltreat Scientific Products, Pepperell, MA, USA) at 5×10^5^ cells/well with RPMI1640 supplemented with 2.5% FBS. The inserts were placed onto wells containing TAM CM or control RPMI1640 medium for THP-1, and incubated for 19 h. The relative number of cells migrated to the bottom chamber was quantified using Celltiter Glo (Promega, Madison, WI, USA). The remaining cells in the inserts were analyzed with Caspase-Glo 3/7 assay (Promega) to confirm that Jurkat cells incubated with TAM CM did not induce apoptosis compared to the control medium.

### Cytokine measurement

The cytokine array was performed using Proteome profiler human cytokine array kit (R&D Systems, Minneapolis, MN, USA) following the manufacturer’s instruction with modifications. For cytokine profiling of polarized TAMs, cytokines were measured in CM collected from THP-1-derived TAM without inhibitor treatments. Cytokines secreted from TAMs treated with BMS-794833 or BMS-5 were analyzed using the supernatant of the polarization culture on day 3 that accumulated secreted factors from TAM into 4T1 tumor CM mixture. Instead of the provided chemiluminescence-based detection, the signals were detected with 800CW-Streptavidin (LI-COR, Lincoln, NE, USA) stained at 1:5000 dilution in the wash buffer. The membranes were scanned using the Odyssey imaging system (LI-COR) and the signal was quantified using Image Studio software (LI-COR). The absolute concentration of cytokines was measured by the Luminex system (R&D Systems).

### Mouse experiment

All animal studies were approved by the IACUC committee of FHCRC. BALB/c and C57BL/6J females were sourced from The Jackson Laboratory. 4T1 cells or Py8119 cells suspended at 1-2 million/150 µl of 33% (v/v) matrigel (Fisher Scientific) were transplanted subcutaneously to the right flank of 6-8 week-old BALB/c or C57BL/6J female mice, respectively. After tumors reached 50-100 mm^3^, mice were randomized into treatment and untreated groups. The mice received intratumoral injection of BMS-794833 at 25 mg/kg dose, BMS-5 at 30 mg/kg dose, or vehicle control twice weekly. BMS-794833 (Selleckchem, Houston, TX, USA) was dissolved into 4% (v/v) DMSO (Sigma-Aldrich), 45% (v/v) PEG300 (Sigma-Aldrich), and 5% (v/v) Tween 80 (Fisher Scientific). BMS-5 (Medchem Express, Monmouth, NJ, USA) was dissolved into 5% DMSO, 30% PEG400, 5% propylene glycol, and 0.5% Tween 80. The tumor size was measured by caliper twice weekly and by weight at the experimental endpoint. The tumor volumes were calculated with the following equation: V= 4/3π×(L/2)×(W/2)×(H/2), (L=length, W=width, H=height). HCI010 model was kindly gifted from Alana Welm (University of Utah) and expanded orthotopically in NSG female mice inbred in-house by Comparative Medicine of FHCRC as described before (DeRose et al., 2013). Py8119 tumors used for CM collection used in THP-1-derived TAM polarization were prepared by injecting Py8119 cells subcutaneously to NSG female mice.

### Flow cytometry analysis and cell sorting

To dissociate the tumor tissue into single cells, tumor tissues up to 1 g were mechanically minced and digested in 2 ml RPMI1640 with 200 µg/ml Liberase TL (Roche by MilliporeSigma), 200 µg/ml Liberase DL (Roche), and 48 unit/ml DNase I (Sigma) at 37°C for 45 min. The dissociated cells were filtered with a 70 µm cell strainer and incubated with red blood cell (RBC) lysis buffer (100 mM Tris, 155.2 mM NH_4_Cl, pH 7.5) for 3 min at room temperature to remove RBCs. The remaining cells were suspended in Flow buffer (1% FBS, 1 mM EDTA, 25 mM HEPES pH 7.0, in PBS) or FACS buffer (5% FBS, 0.05% NaN_3_ in PBS).

To stain the cell surface markers, the cells were first blocked with anti-CD16/32 for 20 min, then stained with fluorescent-conjugated antibodies at 1 µg/ml of each antibody for 20 min. The cells were subsequently stained with a fixable viability dye eFluor 780 (Invitrogen by Thermo Fisher Scientific) for 30 min. The cells for sorting were kept on ice without fixation until analysis. The cells for flow cytometry analysis were fixed for 20 min with a fixation buffer (BioLegend). For staining of intracellular antigens, cells were permeabilized with permeabilization wash buffer (BioLegend) and stained with fluorescent-conjugated antibodies for 20 min. For cytokine analysis, cells were incubated with 5 µg/ml Brefeldin A (BioLegend) in RPMI1640 supplemented with 10% FBS for 4 hours at 37°C before staining. Cells were suspended in 0.1% formaldehyde in PBS and stored at 4°C in the dark until analyses.

The dissociated cells from Py8119 tumors were analyzed by BD FACSymphony and sorted using BD Aria II. The cells from 4T1 tumors were analyzed by BD FACSCelesta and sorted by BD FACSymphony S6. Soring gates of TAM, monocytes, and neutrophils are shown in Figure S3A, C. The gating strategy for 4T1 tumors was applied from (Cassetta et al., 2016). The data were analyzed by FlowJo software. Unstained cells and single stain controls of antibodies with the same fluorophores were used to validate the gating strategy.

For analysis of THP-1-derived TAMs, the THP-1 cells were polarized with 4T1 tumor CM in the presence of BMS-794833 at 7.5 µM or vehicle control for 3 days. The cells were washed with PBS and detached with a cell scraper. THP-1 without induction from suspension culture was washed with PBS and directly used for cell staining. The cells were incubated with Fc blocking reagent on ice for 20 min, stained with fluorophore-conjugated antibodies at 1 µg/ml for 30 min on ice, followed by the fixable viability dye staining for 30 min. The cells were stored in 1% formaldehyde in PBS at 4°C in the dark until analysis using BD FACSCelesta. The antibodies used for flow cytometry and cell sorting are listed in Table S3.

### Histology

At the end of the animal study, tumors and lungs were harvested from mice and fixed in 10% neutral-buffered formalin (MilliporeSigma) for 7 days. The tissues were embedded into paraffin blocks and cut into sections. Hematoxylin-Eosin (H&E) staining of lung sections was performed by the Histopathology core at the FHCRC. The entire tissue was scanned with TissueFAXS (TissueGnostics, Vienna, Austria). The number and area of metastatic foci were evaluated using QuPath software(Bankhead et al., 2017).

For immunohistochemistry of cleaved caspase-3, the tumor sections were deparaffinized, boiled in 10 mM sodium citrate buffer (pH 6.0) for 30 min, and blocked with 2.5% goat serum for 30 min. The sections were stained with cleaved caspase-3 antibody (#9661, Cell Signaling Technology, Danvers, MA, USA) at 1:200 dilution in 1.25% goat serum at 4°C over the weekend. The sections were further incubated with HRP conjugated goat anti-rabbit antibody (ImmPRESS kit, Vector Laboratories, Burlingame, CA, USA) for 30 min at room temperature, and signals were detected by DAB substrate kit (Abcam, Cambridge, UK) followed by hematoxylin staining. After the sections were scanned with TissueFAXS, the area stained with DAB was analyzed with Image J software.

### Isolation and induction of mouse bone marrow-derived cells (BMDCs)

The bone marrow cells were collected from thigh and femur bones from BALB/c or C57BL/6J female mice (The Jackson Laboratory). The bone marrows were flushed with PBS using a syringe, and red blood cells were lysed by incubating in RBC buffer. The cells were incubated in RPMI1640 for THP-1 culture supplemented with human M-CSF at 20 ng/ml (cross-react with mouse, Peprotech, Cranbury, NJ, USA). After 2 days of culture, the medium with floating cells was discarded and replenished with 50% (v/v) 4T1 or Py8119 tumor CM with M-CSF 20 ng/ml. On day 4, M-CSF concentration was raised to 50 ng/ml. Cells were collected for RNA and protein analysis after 7 days from BMDC harvest.

### Quantitative real-time PCR

RNA was extracted using Trizol (Thermo Fisher Scientific) following the manufacturer’s protocol. For RNA extraction from TAM and Ly6C^+^ monocyte cells isolated from tumors, RNeasy Micro Kit (Qiagen, Hilden, Germany) was used following the manufacture’s protocol. cDNA was synthesized with the RT2 HT first strand kit (Qiagen), an qPCR was performed by SYBR green supermix (Bio-Rad, Hercules, CA, USA) with primers purchased from RealTimePrimers.com (Elkins Park, PA, USA). The relative expression levels were calculated with 2-ΔΔCt methods with the internal control as *RPL13A* and normalized to the control sample.

### Immunoblots

For protein lysates for WB and reverse-phase protein array (RPPA), cells were washed twice with PBS and lysed into SDS lysis buffer (2% SDS, 2%, 50 mM Tris-HCl, 5% glycerol, 5 mM EDTA, 1 mM NaF, 1×Protease and Phosphatase Inhibitor Cocktail (Pierce by Thermo Fisher Scientific), 10 mM β-GP, 1 mM PMSF, 1 mM Na_3_VO_4_, 1 mM DTT). The lysates were filtered with 0.22 µm spin-filter. The protein concentration was measured by BCA Protein Assay Kit (Pierce) with bovine serum albumin as a standard. The proteins were denatured with NuPAGE LDS Sample Buffer (Invitrogen) and Reducing Agent (Invitrogen) at 95°C for 5 min. The same protein amount of samples was loaded per lane and separated by SDS-PAGE, and transferred to nitrocellulose membranes with the wet tank method. After blocked with LI-COR Intercept (PBS) Blocking buffer (LI-COR), the membrane was incubated with the primary antibodies at 1:1000 dilution at 4°C overnight, followed by incubation with IRDye 680LT Goat anti-Rabbit IgG (H+L) (LI-COR) or IRDye 800CW Goat anti-Mouse IgG (LI-COR) at 1:5000 for 1 h at room temperature. The signals were detected and quantified with Odyssey infrared imaging system (LI-COR). The primary antibodies are purchased from the following companies; Cell Signaling Technology: P-FAK (#8556), FAK(#3285), P-PYK2 (#3291), P-Paxillin (#2541), Paxillin (#2542), P-Cofilin (#3313), Cofilin (#5175), LIMK1 (#3842), P-SRC family (#2101), FGR (#2755), FYN (#4023), LYN (#2796), P-JAK2 (#3771), JAK2 (#3229), P-STAT3 (#9145), STAT3 (#9139), P-MEK (#9154), MEK1/2 (#9126), P-ERK1/2 (#4377), ERK1/2 (#9102), P-p38 (#9215), p38α (#9217), P-AKT (#4058), AKT (#2920), P-NFκB (#3033), NFκB (#4764); Abcam: PYK2 [YE353] (ab32571); Santa Cruz Biotechnology: c-SRC (sc-19, clone N-16), HCK (sc-72, clone N-30); Sigma-Aldrich: Anti-β-Actin (A1978, clone AC-15).

For RPPA, protein lysates were dot-blotted onto nitrocellulose pad on slide glasses. The slides were washed with 1M Tris-HCl buffer (pH 9.0) for 4-7 days to remove lysate-derived SDS. The protein pads were blocked with LI-COR blocking buffer (PBS), stained with primary antibodies at 1:150 dilution followed by infrared dye-conjugated secondary antibodies at 1:1000 concentration. Spots were analyzed by Array-Pro32 software (Media Cybernetics, Rockville, MD, USA). Signal intensities were normalized to total protein staining by Revert 700 total protein stain (LI-COR).

For tyrosine kinase protein array, Human RTK Phosphorylation Array C1 (AAH-PRTK-1-4, RayBiotech, Peachtree Corners, GA, USA) was used following the manufacturer’s instructions with modification. After incubated with HRP-conjugated antibodies, the protein arrays were incubated with anti-HRP IR800 antibody (LI-COR) and detected by Odyssey imaging system. Gene Ontology enrichment analysis was performed within STRING multiple protein network analysis (Szklarczyk et al., 2019).

### Partial least square regression (PLS-R) analysis

PLS-R analysis was performed using the PLS package of MixOmics package(Rohart et al., 2017) with regression mode, with the number of components as 3. The log_2_-transformed fold change of gene expression data from Figure 3E, the log_2_-transformed fold change of protein phosphorylation data from Figure S4F were used as X variables, and cellular elongation data was used for Y variables.

### Statistics

All the statistical analyses were performed using Graphpad Prism 8.0.1 software. The statistical analyses used for each data are indicated in Figure Legends. *p<0.05, **p<0.01, ***p<0.001, ****p<0.0001. Heatmaps were prepared using pheatmap package in R Studio.

## Results

### Development and characterization of an in vitro TAM polarization model

To develop a physiologically-relevant *in vitro* TAM polarization model, we exposed macrophages differentiated from THP-1 cells, a human monocyte cell line, to tumor CM collected from multiple tumor models (Figure 1A). CM from the mouse triple-negative breast cancer (TNBC) cell line 4T1 induced protrusions in the THP-1-derived macrophages, causing them to exhibit drastically elongated cellular morphology starting from day 2 with peaking at around day 4 (Figure 1B). Given that previous studies have shown that TAM cellular elongation correlates strongly with pro-tumor phenotypes (Benner *et al*., 2019; Cabanel *et al*., 2015; Chen *et al*., 2017; Hollmen et al., 2015; Hu et al., 2021; Porcheray et al., 2005; Su *et al*., 2014; Vereyken et al., 2011), we adopted cellular elongation quantified by live-cell imaging as an indicator of TAM polarization (Figure 1A). We then assessed whether the TAM polarization model reproduces the molecular characteristics of TAMs *in vivo*. Gene expression profiles of macrophages induced by the CM from 4T1 cells upregulated TAM-associated marker genes and temporally induced inflammatory marker genes compared to macrophages cultured without 4T1 CM (Figure 1C). The TAM-related secretory proteins were also detected from the 4T1 CM-induced TAMs (Figure S1A). Consistent with previous reports (Grugan et al., 2012), the TAM model induced both inflammatory (CCL1, Groα, IL-1β, IL-6, IL-8, IL-12, MIP-1α, MIP-1β, CCL5, TNFα, sICAM) and anti-inflammatory cytokines (IL-1RA, IL-4 IL-10, SerpinE1) to 1.4-13.2 and 1.2-1.8 folds, respectively, over control macrophages (Figure S1A). Similar cellular elongation during TAM polarization was observed with CM collected from other breast cancer models (4T1 tumors, Py8119 tumor, a Ras-expressing human mammary epithelial cells, HMLE-Ras, and breast cancer PDX models) (Figure 1D, top), melanoma (B16.F10.Ova), pancreas cancer (Panc02), and colon cancer (MC38 and CT26) (Figure 1D, bottom). In contrast, CM from untransformed HMLE cells did not polarize TAMs. These results provide validation that our tumor CM-induced TAM polarization system is a reliable *in vitro* model for studying TAM function.

**Figure 1.**
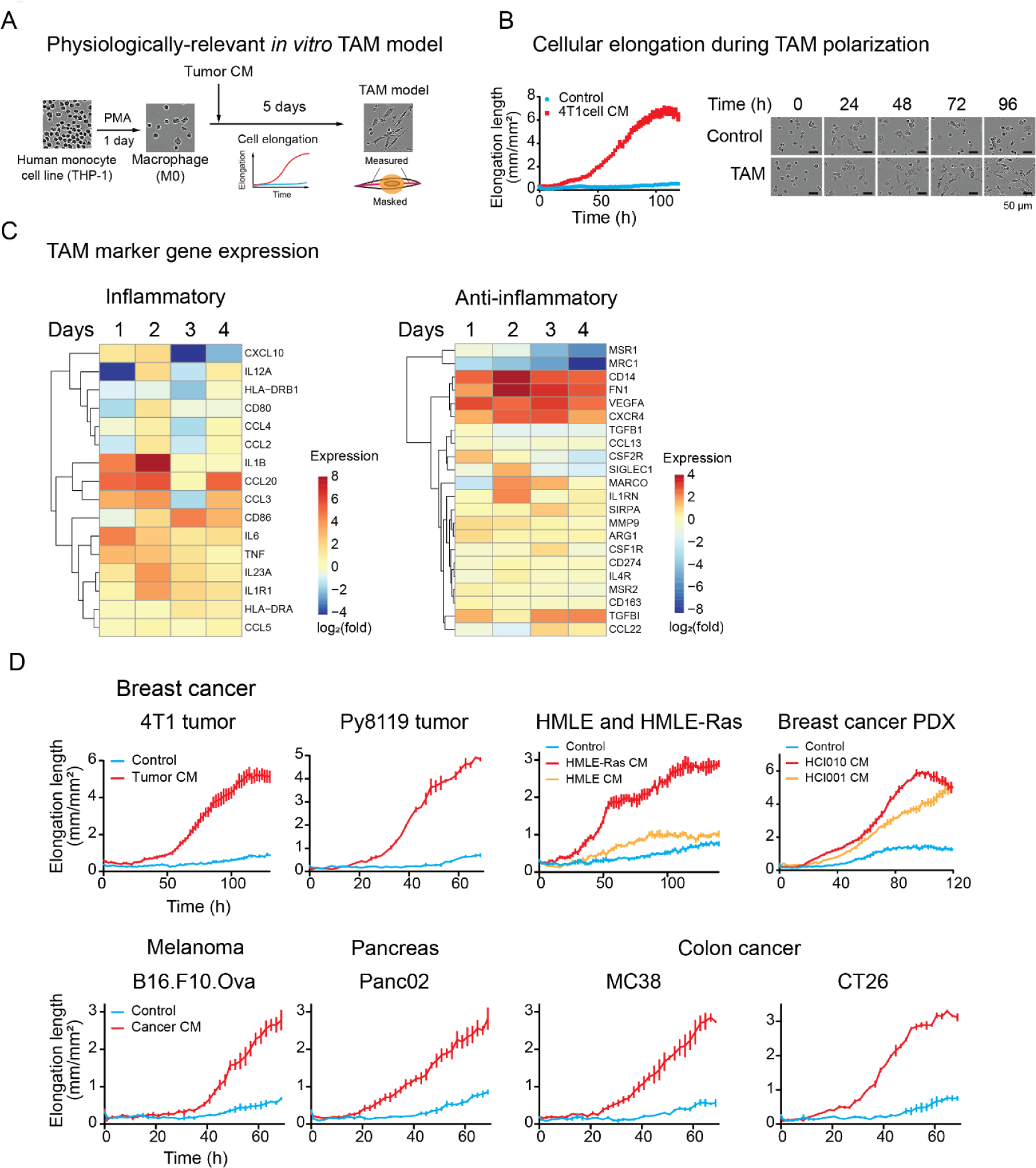
Establishment and characterization of an *in vitro* TAM polarization model. (A) A schematic showing establishment of an *in vitro* TAM polarization model. Human monocyte THP-1 cells were induced by phorbol 12-myristate 13-acetate (PMA) to differentiate into macrophages, followed by culturing in the presence of tumor conditioned medium (CM). Cellular polarization was assessed by cellular elongation measurement via live-cell imaging, as depicted in the illustration. (B) Morphological alteration during TAM polarization in culture. THP-1-derived macrophages were incubated with CM collected from 4T1 cells. Live-cell images were captured every 2 hours under a live cell imaging microscope. The cellular protrusion length per image was measured using an image analysis software. (C) Gene expression of *in vitro* TAM model. THP-1 cells were polarized using CM from 4T1 tumor and collected at the indicated time. The expression of TAM marker genes was analyzed by qPCR. Data are presented as the log_2_ fold change of expression of CM-treated cells over control cells for each day. (D) Validation of CM from multiple tumor models for cellular elongation of THP-1-derived macrophages. (Top) Validation in other breast cancer models. THP-1-derived macrophages were cultured with CM collected from 4T1 tumors, Py8119 tumors, TNBC PDX tumors (HCI010 and HCI001), and human mammary epithelial cell lines, HMLE and HMLE-Ras. CM from HMLE (control) and a cancerous cell line HMLE-Ras, generated by introducing oncogenic Ras gene, were compared in TAM polarization potential. (Bottom) TAM polarization by CM from other cancer types. CMs from mouse melanoma (B16.F10.Ova), pancreatic cancer (Panc02), and colon carcinoma (MC38 and CT26) were used to induce THP-1-derived macrophages. Cellular elongation was measured under live-cell imaging system. The graphs indicate mean ± SEM of measurement at each time point.

### In vitro-derived TAMs promote cancer proliferation and angiogenesis, and inhibits T cell chemotaxis

To further investigate the pro-tumoral behavior of our TAM model and further validate that this model recapitulates in vivo physiology of TAMs, we performed a series of co-culture experiments. Co-culture of 4T1 breast cancer cells with breast cancer CM-induced TAMs stimulated cancer cell proliferation, recapitulating previously-reported growth stimulus effects of TAMs (Figure 2A). Similar cancer cell growth stimulation by TAM was observed with TAMs induced by CM from other cancer models (melanoma, pancreas, and colon cancer) (Figure 2B). To assess pro-angiogenesis ability of TAM, we treated HUVECs with TAM CM that was enriched with secretion factors from polarized TAMs, and monitored the ability of HUVECs forming tube-like structures. Our data show that TAM CM exposure stimulates both length and network branching of endothelial cells (Figure 2C). It is known that TAMs impede T cell infiltration into tumor by secretory factors (Gunderson et al., 2020; Zhang et al., 2018; Zhang et al., 2017). Consistently, TAM CM suppressed Jurkat cell migration in a trans-well migration assay (Figure 2D). Together, these results further validate our *in vitro* TAM polarization model as the system recapitulates *in vivo* ability of TAMs to promote cell proliferation and angiogenesis, and suppress T cell chemotaxis. Therefore, this model offers an opportunity to conduct screening campaigns to identify compounds that re-program pro-tumor TAMs into anti-tumor macrophages.

**Figure 2.**
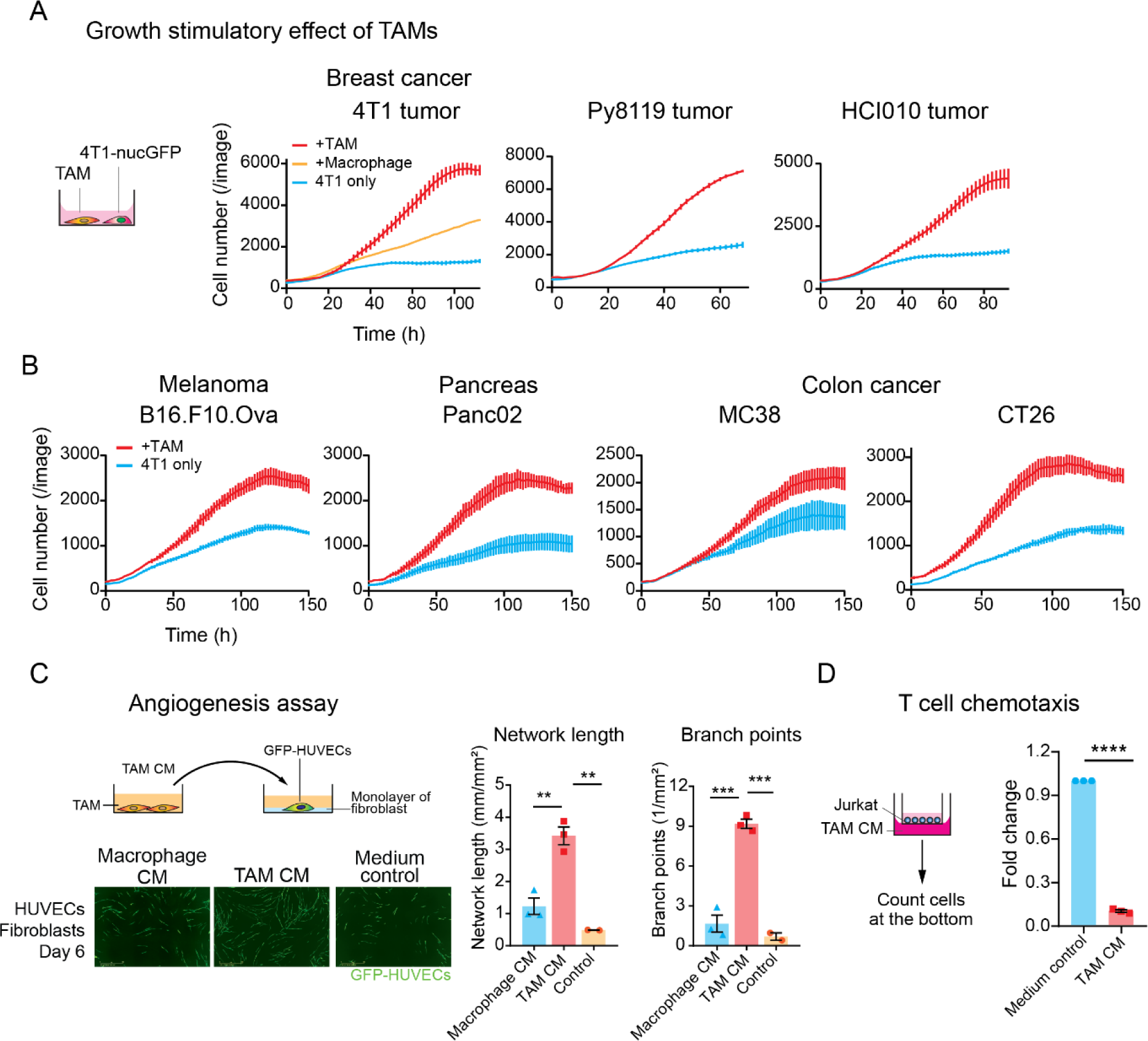
Tumor CM-induced TAMs exhibit pro-tumoral phenotypes. (A) Growth stimulation effect of TAMs on cancer cells. 4T1 cells labeled with nuclear GFP (4T1-nucGFP) were cocultured with breast tumor CM-induced TAMs. THP-1-derived macrophages were polarized with CM collected from indicated breast cancer models (TAMs) or control medium (Macrophages) for 3 days. After TAMs were washed, 4T1-nucGFP cells were seeded on top of TAMs with serum-reduced medium. For control, 4T1-nucGFP was cultured without macrophages (4T1 only). The number of 4T1 cells was counted based on nuclear GFP by image analysis software. The bars indicate mean ± SEM of measurement at each timepoint. (B) Growth stimulation effect of TAMs induced by CM from melanoma (B16.F10.Ova), pancreatic cancer (Panc02), and colon cancer cell lines (MC38 and CT26). The experimental details are the same as (A). (C) Stimulation of tube formation of HUVECs by TAMs. GFP-expressing HUVECs were seeded on a monolayer of fibroblasts. HUVECs were cultured with TAM CM collected from TAM induced by 4T1 tumor CM. (Left) Wide-field image obtained after 6 days of incubation. (Right) Network length and branch points of GFP-HUVEC networks on day 6 by live cell imaging analysis. The bar graphs show mean ± SEM of n=3 (macrophage CM, TAM CM), n=2 (medium control). **p<0.01, *** p<0.001, One-way ANOVA with Tukey’s multiple comparison tests. (D) T cell chemotaxis assay by TAM model. Jurkat cells seeded in culture inserts were incubated with TAM CM collected from TAM induced with HCI010 CM. The relative number of Jurkat cells migrated to the bottom was quantified by luminescent-based cell viability assay. Individual experimental values are shown as dots (n=3). **** p<0.0001, One-way ANOVA with Tukey’s multiple comparison tests.

**Figure 3.**
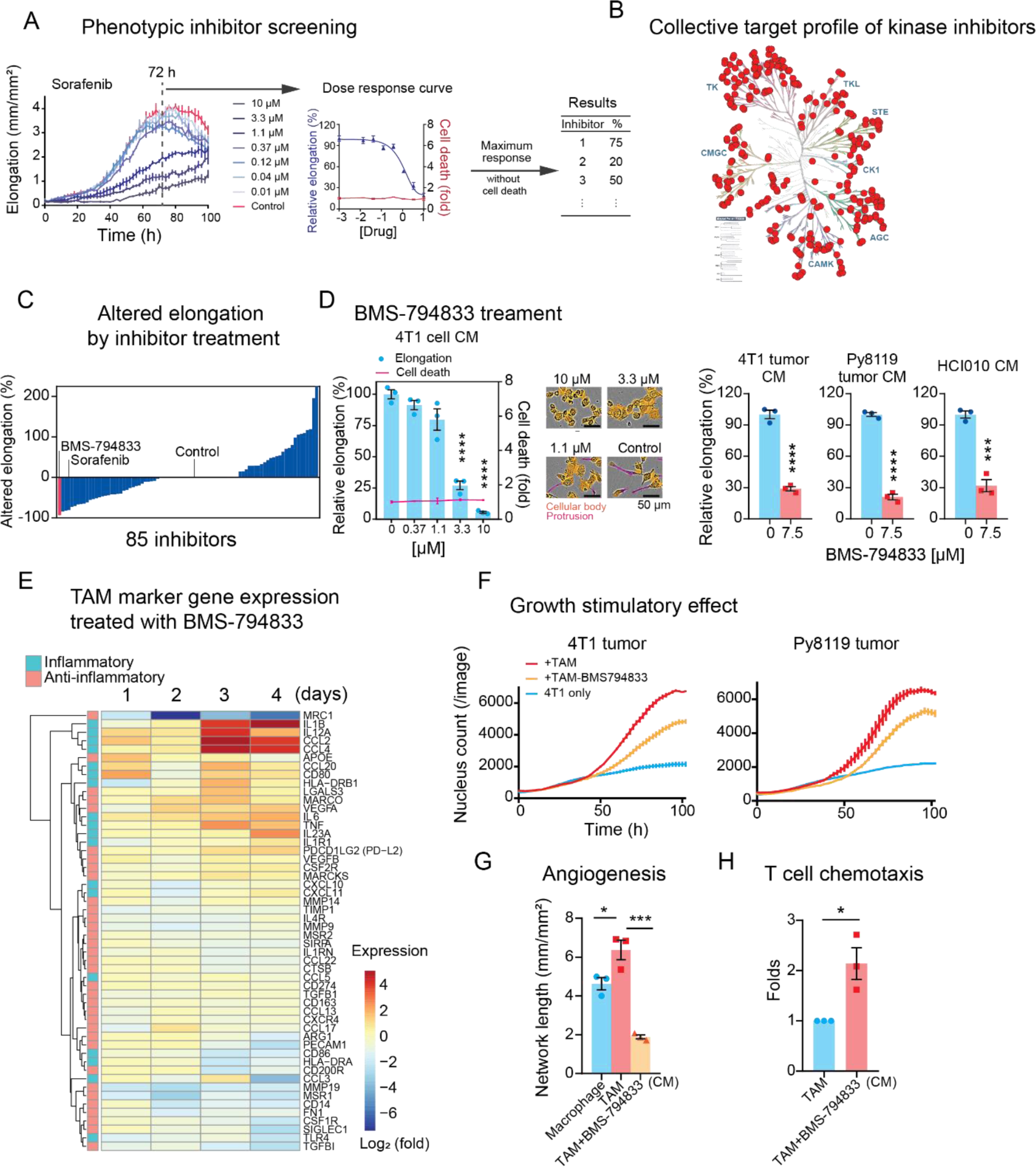
Kinase inhibitor screening in the *in vitro* TAM polarization model identified BMS-79833 as a potent TAM polarization inhibitor. (A) A schematic showing the screening procedure for 85 kinase inhibitors in an *in vitro* TAM polarization model. Response to sorafenib is shown as a representative. TAMs were polarized under eight serial doses of kinase inhibitors ranging from 0 to 10 µM. Relative cellular elongation levels of inhibitor-treated TAMs compared to control were measured as the effect of inhibitors. A red fluorescent cell death indicator dye, YOYO-3, which detects disrupted cellular membranes, was supplemented to detect cellular deaths caused by kinase inhibitors. (B) Collective target kinase profiles of the 85 screened kinase inhibitors. The selected kinase inhibitors cover 289 kinases from various kinase families, with lower than 50% residual activities at 0.5 µM. The kinase tree was prepared using KinMap (Eid et al., 2017). (C) Changes in cellular elongation in response to inhibitor treatment. Inhibitors are plotted based on the maximum altered elongation levels at up to 10 µM dose. Inhibitors causing cellular death were regarded as causing no change in elongation. (D) Inhibitory effect of BMS-794833 on cellular elongation of THP-1-derived TAM polarization. (Left) Elongation of THP-1-derived TAMs treated with a serial dose of BMS-794833 in the presence of 4T1 cell CM. The relative cellular death of inhibitor-treated TAMs was overlayed on the graph. The bars and plots are the means with ± SEM of n=3. ****p<0.0001, One-way ANOVA with Dunnett’s multiple comparison test. (Middle) Representative images of THP-1-derived TAM with BMS-794833 at the indicated concentration at 72 h from polarization. Cell protrusions (measured as cellular elongation) and cellular body masked are labeled with magenta and orange, respectively. (Right) TAM elongation with CM from 4T1 tumor, Py8119 tumor, and breast PDX HCI010 with or without BMS-794833 at 7.5 µM. ****p<0.0001, unpaired two-tailed Student’s t-test. (E) Time-course analyses of TAM marker gene expression of BMS-794833-treated TAMs. Log_2_-transformed fold changes of gene expression of BMS-794833-treated TAMs at 7.5 µM compared to control TAMs were shown in heatmaps. Genes with inflammatory or anti-inflammatory properties are shown in the left color panels. (F) Growth stimulation effect of TAM treated with BMS-794833. 4T1-nucGFP were cocultured with BMS-794833-treated TAMs at 7.5 µM induced by 4T1 tumor CM or Py8119 tumor CM under a reduced serum condition. The growth of 4T1 was evaluated by nuclear GFP counts. The bars indicate mean ± SEM of measurement at each timepoint. (G) Tube formation ability of TAM treated with BMS-794833. CM was collected from TAM with or without BMS-794833 treatment at 7.5 µM induced by 3 independent batches of HCI010 CM. The TAM CM was supplemented to tube formation assay of GFP-expressing HUVECs grown on fibroblast layer. *p<0.05, ***p<0.001, one-way ANOVA followed by Tukey’s multiple comparisons. (H) T cell chemotaxis assay. Jurkat cells migrated through the bottom of the chamber with CM from TAMs polarized with BMS-794833 at 7.5 µM. The relative number of migrated cells was evaluated by luminescent-based cell assay. CM from TAM induced by three independent batches of HCI010 CM were used for this study. *p<0.05, unpaired two-tailed Student’s t-test.

### Kinase inhibitor screening on the in vitro TAM model identified BMS-794833 as a potent inhibitor of TAM polarization

Protein kinases (kinases) are critical components of cellular signaling networks that transmit extra/intracellular stimuli to cellular responses. Given that TAM polarization is an external stimuli-regulated process, we hypothesized that kinases will play a major role in this process, and therefore, kinase inhibitors may represent useful pharmacological agents to inhibit pro-tumor TAM polarization. To examine this further, we used our model system and cellular elongation as an indicator of TAM polarization levels to screen a curated collection of 85 kinase inhibitors that achieves broad kinome coverage (289 kinases out of 298 kinases measured), with lower than 50% residual activities at 0.5 µM (Figure 3B). We tested this inhibitor collection at eight serially-diluted concentrations (0 - 10 µM) on our TAM model using THP-1 cells induced with CM from 4T1 cultured cells (Figure 3A). The efficacy of each inhibitor on TAM polarization was quantified as inhibition of cellular elongation compared to vehicle control. To omit inhibitors causing cellular death, we supplemented the assay with YOYO-3, a red fluorescent dye that stains membrane compromised cells. The response to each inhibitor was evaluated at the highest concentration ranging from 0 to 10 µM that did not exhibit cellular death. As a result, of the 85 tested inhibitors, 33 inhibitors caused a reduction in elongation by 3-93%, 26 inhibitors showed no change and/or induced cellular death, and surprisingly, 26 caused an increase in elongation by 14-225% (Figure 3C, Table S1). Sorafenib, a multikinase inhibitor of Raf, VEGFR, and PDGFR, has been previously shown to inhibit M2 (anti-inflammatory) macrophage phenotype and restore inflammatory cytokine expression (Deng et al., 2016; Edwards and Emens, 2010; Sprinzl et al., 2015). In our assay, sorafenib efficiently suppressed cellular elongation at 3.3 µM or higher, with 81.5% inhibition at 10 µM (Figure 3A) and was the fourth-most effective inhibitor (Figure 3C), validating the reliability of the screening system.

Among the tested inhibitors, BMS-7948333, a dual inhibitor of c-MET and VEGFR2(J.Fargnoli et al., 2010), was the most potent inhibitor of cellular elongation during TAM polarization, with 93% inhibition at 10 µM with no evidence of cell death (Figure 3D, Figure S1B). We confirmed the inhibitory effect of BMS-794833 on cellular elongation in our other TAM polarization models using breast cancer CM (4T1 tumor, Py8119 tumor, and PDX) as well as CM from other cancer models (Figure 3D, Figure S1C). Time-course transcriptional analyses of BMS-794833-treated cells showed a marked decrease in expression of the TAM marker (MRC1/CD206), and enhanced expressions of inflammatory cytokine-coding genes, such as *IL1B*, *CCL2*, *CCL4*, *IL12A*, *IL6*, and *TNFA* (Figure 3E). Cytokines secreted from BMS-794833-treated TAMs were also measured using protein arrays and absolute quantification using Luminex (Figure S1D). BMS-794833 treatment suppressed secretion of anti-inflammatory cytokines IL-4, IL-13, and IL-10, and enhanced inflammatory cytokines such as IL-6, IL-8, IL-27, MIP-1α/β, CCL2, TNFα, and IL-1β. These results demonstrate that BMS-794833 could abrogate TAM polarization and re-program TAMs to proinflammatory phenotypes.

### BMS-794833 inhibits TAM polarization and pro-tumoral phenotypes

We next investigated the effect of BMS-794833 on the protumoral activities of TAMs. BMS-7949833 treatment decreased the growth stimulatory effect on 4T1 cells following coculture with 4T1 or Py8119 tumor CM-induced TAMs (Figure 3F), as well as other cancer models (Figure S1E). Additionally, CM collected from BMS-794833-treated TAMs lost angiogenesis stimulation potential of TAMs, and restored T cell chemotaxis ability (Figure 3G, H). We further validated the inhibitory effect of BMS-794833 on TAM polarization models prepared using bone marrow-derived cells (BMDCs) from BALB/c and C57BL/6J mice induced with CM from 4T1 and Py8119 tumors, respectively. In agreement with our previous results, BMDC-derived TAMs exhibit elongated morphology and induced expression of TAM marker genes, and BMS-794833 treatment suppressed both cellular elongation and marker gene expression (Figures S1F, G). Taken together, these results demonstrate that BMS-794833 identified through the phenotypic screen can impede TAMs from enacting their protumoral functions *in vitro* and re-program them into pro-inflammatory phenotype.

### BMS-794833 treatment suppressed breast tumor growth

We next investigated the effect of BMS-794833 on primary breast tumor growth in mice. First, we evaluated the effect of BMS-794833 on the proliferation of breast cancer cells in monoculture. BMS-794833 inhibited cellular growth of 4T1 breast cancer cells by 28% at 10 µM (Figure 4A). Py8119 cells were slightly more sensitive, with IC_50_ at 4.1 µM and inhibition by 87% at 10 µM. These results showed that BMS-794833 has a modest inhibitory effect on breast cancer cell growth. Next, we administered BMS-794833 to 4T1 tumor transplanted subcutaneously in BALB/c mice. Biweekly treatment of BMS-794833 at 25 mg/kg dose intratumorally significantly reduced tumor weight by 52% compared to vehicle control (Figure 4B). Reduction of tumor growth was also observed in Py8119 tumors grown subcutaneously in C57BL/6J mice (Figure 4C). Immunohistochemistry of cleaved caspase-3 in BMS-794833-treated tumors revealed that BMS-794833-treatment caused an increased trend of apoptosis (Figure S2A). The BMS-794833 treatment also decreased lung metastasis in the 4T1 tumor model (Figure 4D). Py8119-bearing mice did not develop visible lung metastatic foci in either control or treated tumors within the experimental period (data not shown). These results demonstrate that BMS-794833 suppresses tumor growth and metastasis *in vivo*.

**Figure 4.**
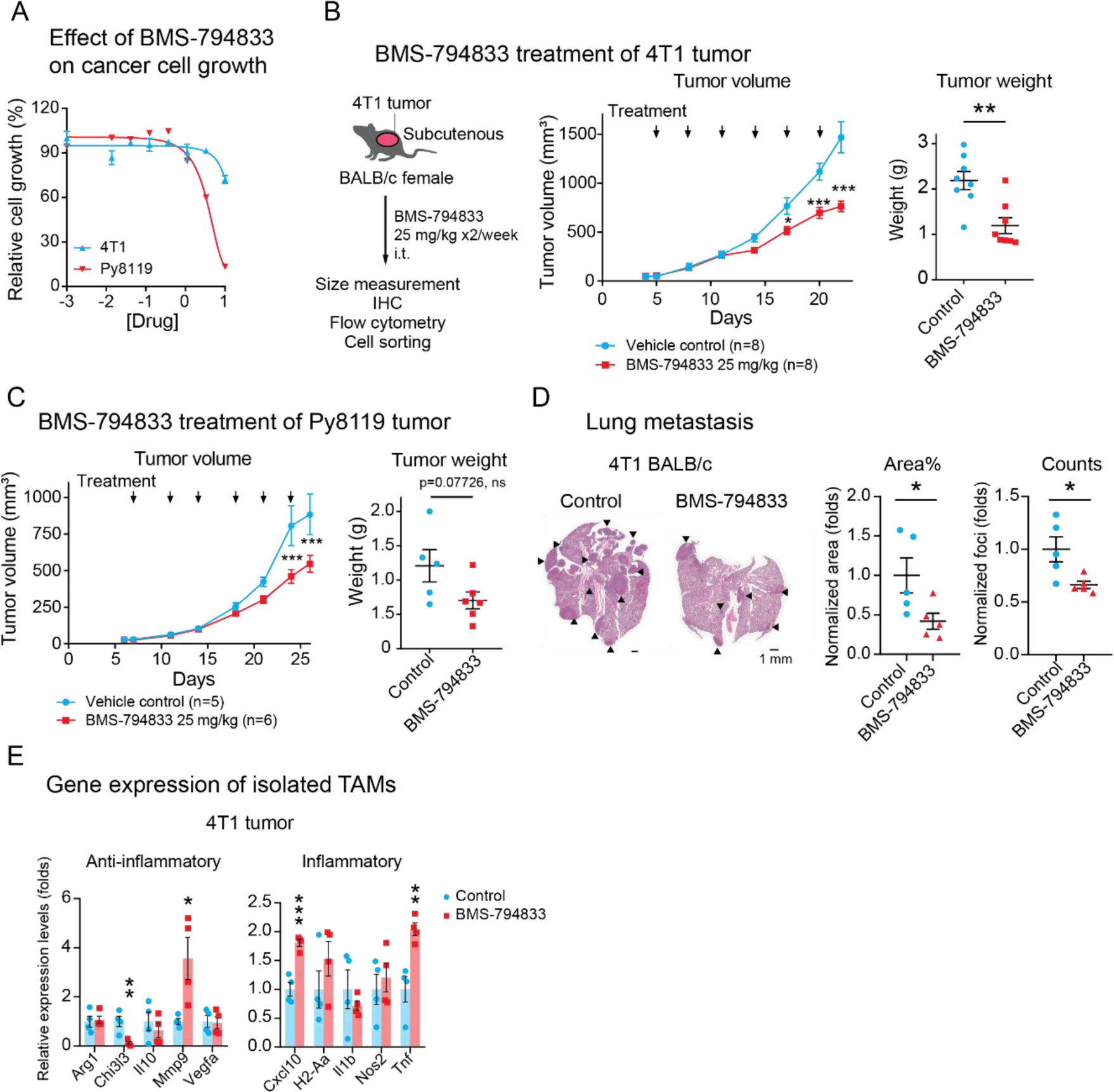
BMS-794833 treatment suppressed breast tumor growth. (A) Effect of BMS-794833 on the proliferation of TNBC cell lines, 4T1 and Py8119 cultured in a plate. 4T1-nucGFP or Py8119 were cultured under the presence of a serial dose of BMS-794833. Cell proliferation of 4T1 was evaluated by nuclear GFP count, and of Py8119 by cell confluency. N=3, with mean ± SEM. (B) Tumor growth of 4T1 tumors treated with BMS-794833. (Left) A schematic of experimental design. BALB/c female mice with subcutaneous 4T1 tumor were treated biweekly with either BMS-794833 (25 mg/kg) or vehicle control with intratumoral injections. (Middle) Growth curves of 4T1 tumors. The timing of drug administration is indicated with black arrows. **p<0.01, multiple t-test at each timepoint with Holm-Sidak correction (α<0.05), n=8. (Right) Tumor weight at the experimental endpoint. Mean ± SEM. **p<0.01, unpaired unpaired two-tailed Student’s t-test. (C) Tumor growth of Py8119 tumors treated with BMS-794833. (Left) Growth curves of Py8119 tumors. CB57BL/6J female mice with subcutaneous Py8119 tumor were treated biweekly with either BMS-794833 (25 mg/kg) or vehicle control with intratumoral injections. The timing of drug administrations is indicated with black arrows. ***p<0.001, multiple t-test at each timepoint with Holm-Sidak correction (α<0.05), n=5 (control), n=6 (treatment). (Right) Tumor weight at the experimental endpoint. Mean ± SEM. Statistical significance was calculated using unpaired two-tailed Student’s t-test. (D) (Left) Representative images of H&E staining of lung sections from BMS-794833-treated or vehicle-treated 4T1 tumor-bearing mice. Black arrows indicate metastatic tumors. (Middle) Normalized percentage of the tumor area and (right) the normalized number of metastatic foci in the lung sections. The lines indicate the mean ± SEM. *p<0.05, unpaired two-tailed Student’s t-test. (E) Gene expression of TAMs (CD45^+^CD11b^+^Ly6C^-^Ly6G^-^F4/80^+^) isolated from 4T1 tumor treated with BMS-794833. Expression of inflammatory and anti-inflammatory genes in the sorted TAM population (the gate is shown in Figures S2C, D) was measured by qPCR. p<0.05, **p<0.01, **p<0.001, unpaired two-tailed Student’s t-test.

### BMS-794833 abrogates TAM polarization to retain inflammatory phenotypes

To investigate whether BMS-794833 treatment impacts the profiles and characteristics of TAMs and other immune cells, we analyzed 4T1, and Py8119 tumors treated or untreated with BMS-794833 by flow cytometry (Figure S3). BMS-794833 caused a significant decrease in live cells (Figure S3B), consistent with cleaved caspase-3 staining observed in tumor sections (Figure S2A). In both tumor models, BMS-794833 treatment did not change the percentage of overall immune populations (CD45^+^ cells) (Figure S3C), myeloid population, and most marker expressions (Figures S3D-H), and most lymphoid populations (Figure S3I). TAM population (CD45^+^CD11b^+^Ly6G^-^Ly6C^-^F4/80^+^) exhibited a slight decrease under treatment but was not statistically significant (Figure S3E). The most prominent change in population was in monocytes, defined by Ly6C immature marker, which was significantly increased in 4T1 and less significantly in the Py8119 model (Figure S3E). Blood-derived monocytes are a major source of TAMs, continuously replenishing TAM in tumor tissues (Qian et al., 2011). An increase in monocytes may be due to either increased chemoattractants, such as CCL2 and MIP-1α/β increased in THP-1-derived TAM by BMS-794833 treatment (Figures 3E, S1D), or inhibition of polarization into TAMs. To investigate whether BMS-794833 causes incomplete TAM polarization, surface markers of THP-1-derived TAMs treated or untreated with BMS-7948333 were analyzed. Untreated TAMs exhibited increased expression of CD11b (a myeloid cell marker), CD68 (a pan-macrophage marker), and CD14 (monocytes/macrophage marker) compared to the parental THP-1 cells (Figure S2B). BMS-794833 treated TAMs retained a similar level of CD11b^+^ cells, decreased cells expressing CD68 and CD14 at the cell surface. These results suggest that the BMS-794833 has inhibitory roles on complete differentiation/polarization of monocytes to TAM.

Next, we investigated gene expression of TAMs and their precursor cells, monocytes, in BMS-794833-treated tumors. The TAM and Ly6C^+^ monocyte populations of 4T1 and Py8119 tumors were isolated through FACS (Figures S2C, D). Gene expression analyses of TAMs isolated from BMS-794833-treated 4T1 tumors significantly upregulated inflammatory marker genes (Cxcl10, Tnf), and suppressed protumoral TAM marker gene (Chi3l3) (Figure 4E). Ly6C^+^ monocytes from BMS-794833-treated 4T1 tumors exhibited a similar decrease in the anti-inflammatory Chi3l3 and increase in inflammatory markers, but to a lesser extent (Figure S2E). In the Py8119 tumors, the gene expression of TAM and Ly6C^+^ monocytes were less significant; neither TAM nor monocytes of Py8119 tumor induce TNFα, but notably, TAM increased Nos expression (Figure S2E). These results demonstrate that BMS-794833 induced inflammatory phenotypes in intratumoral TAMs and Ly6C^+^ monocytes.

### BMS-794833 suppresses TAM polarization through polypharmacological effect

We sought to investigate the mechanism by which BMS-794833 impairs TAM polarization. Although c-MET and VEGFR2 are considered to be two main targets of BMS-794833, *in vitro* target profiling revealed that BMS-794833 inhibited 29 kinases to below 50% residual activity at 0.5 µM (Figure 5A, Figure S4A, Table S2). To examine whether c-MET and VEGFR2 are responsible for TAM polarization, we first tested selective inhibitors for c-MET and VEGFR2, JNJ-38877605 and cediranib, respectively, on the TAM model. Neither compound inhibited TAM elongation (Figure S4B), suggesting that the inhibitory effect of BMS-794833 results from inhibition of either another single target or a polypharmacological effect on multiple targets. To test this further, we also inhibited the top 18 kinases targeted by BMS-794833 using selective inhibitors, but a majority of these inhibitors were not as potent as BMS-794833 (Figure 5B). Only inhibitors chemically-analogous to BMS-794833 showed a similar level (cabozantinib) or less (BMS-777607) inhibition on cellular elongation during TAM polarization (Figure S4C). These results suggest that targeting a single kinase is insufficient to suppress TAM elongation, and that inhibition of multiple signaling pathways is required to effectively block TAM polarization.

**Figure 5.**
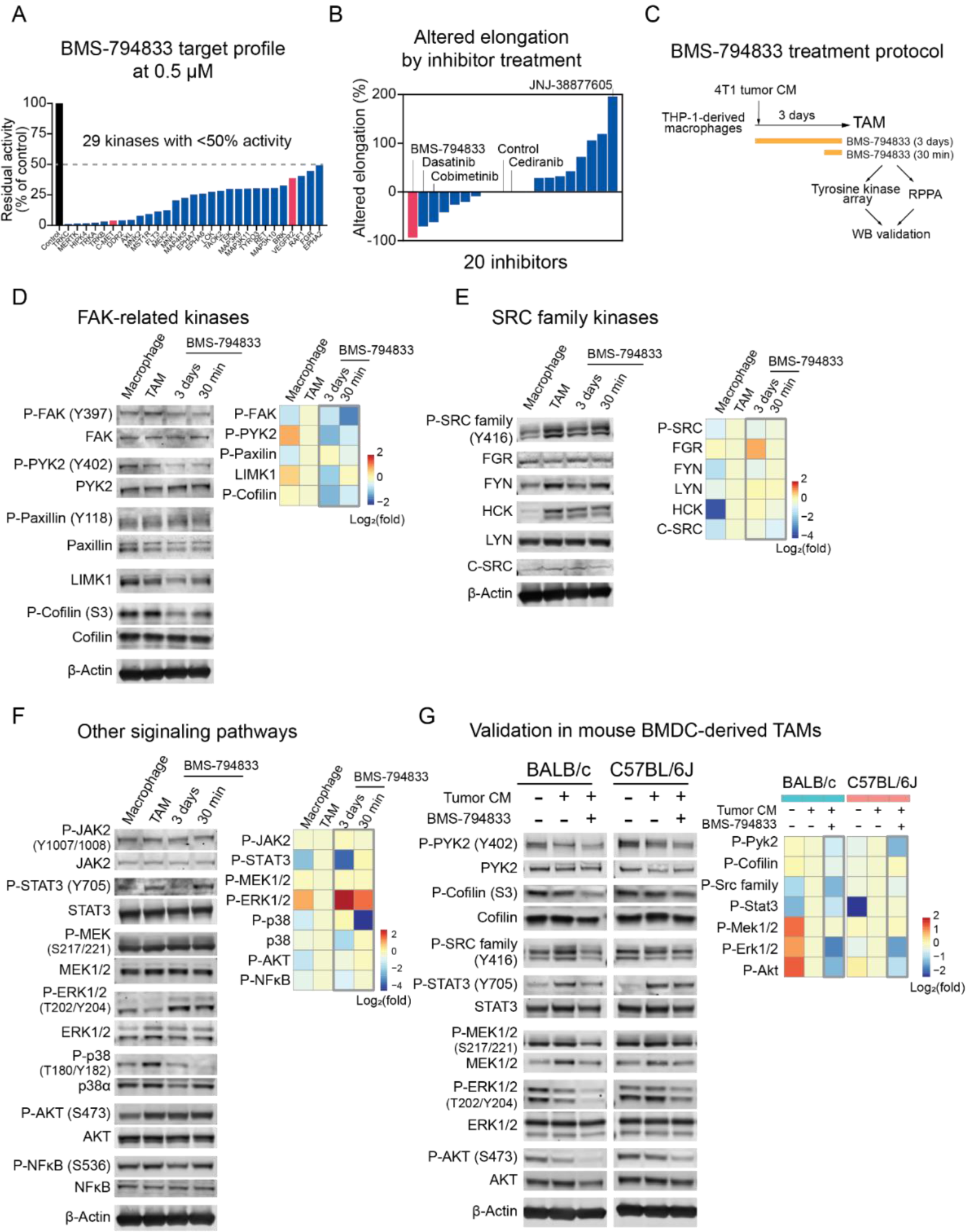
BMS-794833 targets multiple signaling pathways in TAMs. (A) Target kinase profile of BMS-794833. Residual activities of kinases under the presence of BMS-794833 at 0.5 µM were evaluated in *in vitro* acellular system with synthetic substrates (Rata *et al*., 2020). Also, see Figure S4A and Table S2. (B) Changes in cellular elongation caused by inhibitors targeting the top BMS-794833-targeted 18 kinases to below 30% at 0.5 µM. A total of 20 inhibitors, including BMS-794833 as a comparison and a VEGFR2 inhibitor cediranib are shown. Inhibitors are plotted based on the maximum altered elongation observed at below 10 µM dose. Inhibitors that caused cell death were regarded as no change in elongation. (C) Experimental design for protein analysis of BMS-794833-treated TAM. Macrophages prepared from THP-1 were polarized with 4T1 tumor CM for 3 days. BMS-794833 was supplemented to the culture at 7.5 µM final concentration simultaneously with CM addition, or 30 min before sample collection on day 3. The resulting TAMs were analyzed with protein arrays followed by validation by western blotting. (D) Phosphorylation of FAK and cytoskeleton-related proteins. (Left) Representative images of the blot showing FAK and actin-regulation-related proteins. (Right) Heatmap showing quantification of signal intensities. The log_2_ fold change of normalized signal intensity to either total protein or β-actin (LIMK1) is shown. (E) Phosphorylation of SRC family kinases. (Left) Representative images of the blot showing phosphorylated and individual SRC family kinases. The antibody for phosphorylated SRC family kinases detects all SRC family kinases. (Right) Heatmap showing quantification of signal intensities. The log_2_ fold change of normalized signal intensity to β-actin is shown. (F) Immunoblot analyses of additional signaling pathways. (Left) Representative images of the blot showing STAT3, MAPKs, AKT, and NFκB. (Right) Heatmap showing quantification of signal intensities. The log_2_ fold change of normalized signal intensity to either total protein or β-actin is shown. (G) Immunoblot images of mouse BMDC-derived *in vitro* TAM model cells. BALB/c or C57BL/6J-derived BMDC were induced with M-CSF for 2 days, followed by 4T1 (BALB/c) or Py8119 (C57BL/6J) tumor CM supplemented with M-CSF for 5 days, with or without BMS-794833 at 7.5 µM. (Left) Representative images of the blot. (Right) The log_2_ fold change of normalized signal intensity to either total protein is shown.

### Inhibitory effect of BMS-794833 on time-dependent signal activation during TAM polarization

To characterize the effect of BMS-794833 further, we performed a phosphorylated tyrosine kinase array (Figure S4D, C) and reverse-phase protein array (RPPA) screening followed by WB validation (Figure 5C). TAMs polarized with 4T1 CM were treated with BMS-794833 for full-time (3 days) or the last 30 min of the polarization period. BMS-794833 treatment decreased phosphorylation of proteins related to focal adhesion-cytoskeleton regulation, SRC family kinases, JAK-STAT3, p38 MAP kinase, AKT, and NF-κB (Figure 5D, E, F). The results also revealed differential response at the time of inhibition by BMS-794833; phosphorylation of FAK, PYK, cofilin, and p38 MAPK can be inhibited at the last 30 minutes of the polarization period, whereas total LIMK1, phosphorylated STAT3, AKT, and NFκB were downregulated only upon full-time treatment. In detail, phosphorylated FAK, PYK2, and cofilin decreased within 30 min and remained low after 3 days of BMS-794833 treatment, while the total protein level of LIMK1, an upstream kinase that phosphorylates cofilin (Yang et al., 1998), decreased after 3 days (Figure 5D). Among the 9 SRC family kinases (Kim et al., 2009), THP-1-derived TAMs expressed HCK, FGR, LYN, FYN, and a low amount of c-SRC (Figure 5E). Phosphorylation of total SRC family kinases was induced in TAM polarization and decreased by 3-days BMS-794833 treatment (Figure 5E). Phosphorylated STAT3, a transcription regulator of immunosuppressive cytokines, MMPs, and angiogenesis factors (Irey *et al*., 2019) was decreased significantly after long-term BMS-794833 treatment (Figure 5F). The phosphorylation level of p38 MAPK, a conditional regulator of pro- or anti-inflammatory response (Raza et al., 2017), dropped within 30 min and also decreased total protein amount after 3 days. Phosphorylated ERK1/2 was increased by BMS-794833 treatment within 30 min and remain high on 3 days (Figure 5F). However, in the validation experiment with mouse BMDC-derived TAMs, phosphorylation of Erk1/2 and Mek1/2 decreased, indicating that the increase of phosphorylated ERK is a THP-1-specific observation (Figure 5G). Downregulation of phosphorylated Src family, Pyk2, Cofilin, Stat3, and Akt by BMS-794833 were validated in the mouse BMDC-derived TAM model (Figure 5G).

As TAMs dynamically regulate signaling activation over time, we further investigated time-dependent activation of signaling pathways during TAM polarization, leveraging the advantage of our *in vitro* system (Figure S4F). We observed an increase in the phosphorylation of AKT, ERK1/2, and p38 MAPK within one hour in response to 4T1 tumor CM, followed by an increase in the phosphorylation of NF-κB, which peaked at 24 hours, indicating activation of these proteins is an early event in TAM polarization. In contrast, phosphorylation of FAK, cofilin, SRC family, and STAT3 was increased from day 2, indicating that these proteins are activated in a late event. Importantly, BMS-794833 treatment suppressed induction of p38 MAPK, NF-κB, FAK, Cofilin, SRC family, and STAT3, suggesting that BMS-794833 targets both early and late-stage protein signaling during TAM polarization. Together, these data highlight temporal activation of multiple pathways during TAM polarization, and the effect of BMS-794833 on these signal cascades.

### Systems-based analysis of TAM polarization and BMS-794833 treatment

TAMs exhibit dynamic changes in gene expression and protein phosphorylation during the polarization period (Figures 1, 3, 5). To evaluate relative contributions of each change to TAM polarization and to the response to BMS-794833, we performed partial least square regression (PLS-R) analysis of time-course gene expression and protein phosphorylation of TAMs treated or untreated with BMS-794833, with cellular elongation as a response (Figure S5). Component 1 (40% explained variance) delineated time-dependent TAM polarization, and component 2 (19% explained variance) discriminated BMS-794833 treated and untreated groups (Figure S5A). The top 30 standardized coefficient of component 2 includes positive coefficients (towards BMS-794833 treated group) of inflammatory cytokine-coding genes and negative coefficients (towards untreated group) of phosphorylated STAT3, cofilin, p38 MAPK, and total LIMK1 (Figure S5B, C). These results indicate that the effect of BMS-794833 can be explained by both increases in inflammatory cytokine expression and a decrease in phosphorylated/total proteins involve in TAM polarization.

### Targeting multiple signaling pathways is necessary for abrogating TAM polarization

To investigate the contribution of the individual signaling pathway in TAM polarization, we tested selective inhibitors for FAK, LIMK1, SRC family, and STAT3 (Figure 6A). Two potent FAK inhibitors (PF00562271 and NVP-TAE226) showed an inconsistent effect on TAM elongation and did not provide conclusive evidence for the importance of FAK. A highly-selective LIMK inhibitor BMS-5 inhibited cell elongation, consistent with the role of LIMK1 in actin polymerization. LIMK1 is predominantly expressed in the TAM population of breast cancer patient tumors (Figure S6A). BMS-5 treatment upregulated secretion of inflammatory cytokines (Figure S6B) and lowered the growth stimulation effect of TAMs (Figure S6C). However, BMS-5 did not show any tumor-suppressive effect on 4T1 tumor-bearing mice (Figure S6D). Western blot analyses revealed that BMS-5 suppresses phosphorylation of FAK, cofilin, and SRC family kinases, but phosphorylation of STAT3 remained unchanged (Figure 6B). A STAT3 inhibitor, Stattic (Schust et al., 2006), had a modest effect on TAM elongation at 10 µM (Figure 6A). The immunoblot confirmed suppressed phosphorylation of STAT3 (Figure 6B), and also showed decreased phosphorylated SRC. However, phosphorylation of FAK, cofilin and total LIMK remained unchanged. Src inhibitor-1, which inhibits most SRC family kinase, stimulated cellular elongation (Figure 6A). Immunoblot showed that Src inhibitor-1 induces phosphorylated SRC, FAK, and STAT3 within 30 min, and increased LIMK1 expression after 3 days (Figure 6B). The results indicate that SRC signal blockade results in compensatory induction of SRC and other TAM-related signals.

**Figure 6.**
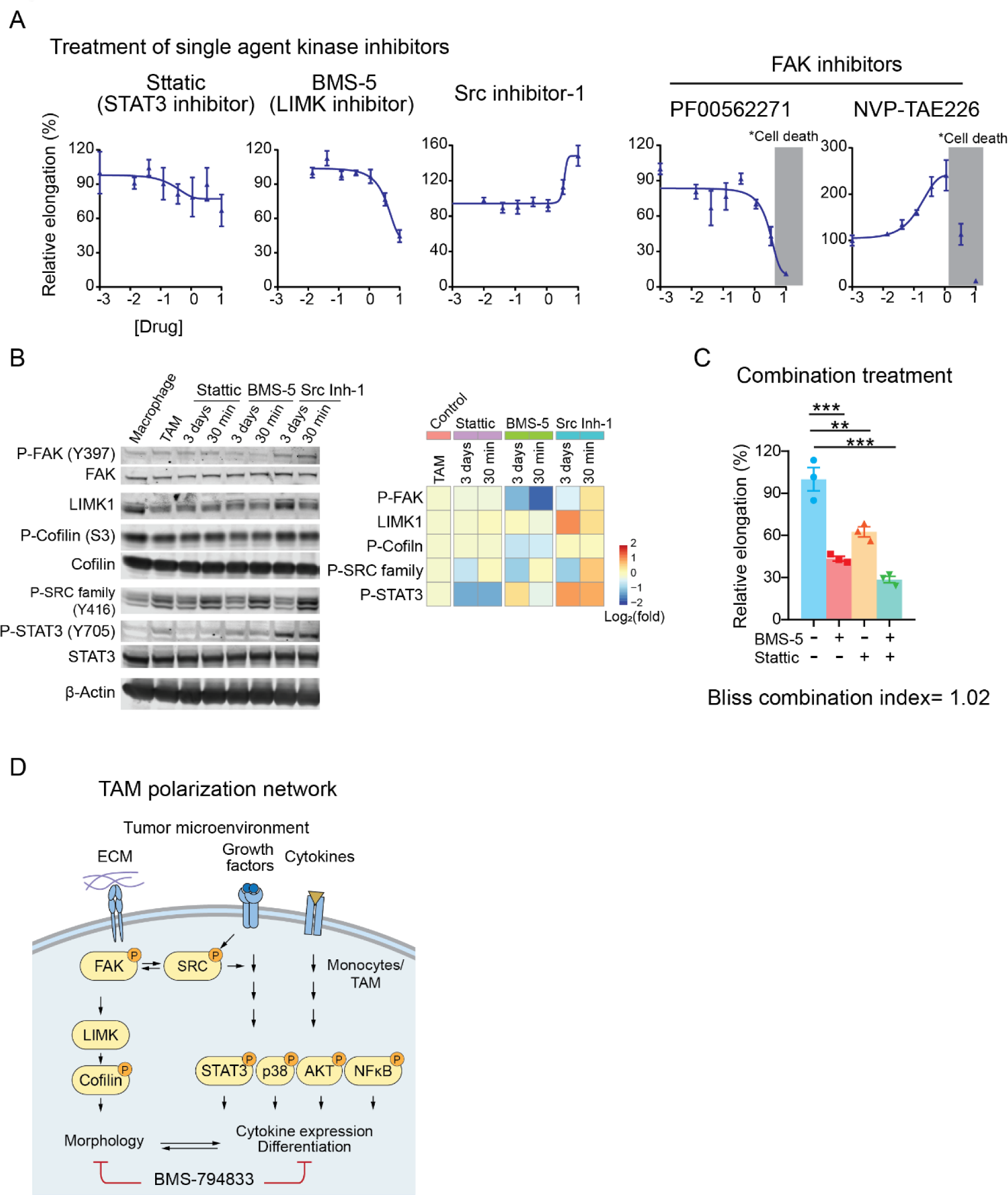
Targeting multiple signaling cascades is required for impeding TAM polarization. (A) Dose-response curve of selective inhibitors for STAT3 (Stattic), LIMK1 (BMS-5), SRC family kinases (Src inhibitor-1), and FAK (PF-00562271 and NVP-TAE226) on TAM polarization using 4T1 tumor CM. Inhibitors were tested within 0 to 10 µM range and dose response was plotted in log_10_ scale. The concentration causing cellular death is shaded with gray. The error bars depict the mean ± SEM of 3 replicates. (B) Immunoblots of TAM polarized in the presence of Stattic, BMS-5, and Src Inhibitor-1 (Src Inh-1). Inhibitors were administered to TAM polarization culture at the time of 4T1 tumor CM induction starts (3 days), or the last 30 min before sample collection at day 3 (30 min) at 10 µM concentration. (C) Combination treatment of Stattic and BMS-5. BMS-5 (5 µM) and Stattic (5 µM) were supplemented as either a single agent or in combination to TAM polarization culture with 4T1 tumor CM. The cellular elongation after 120 h culture is shown. The bar graphs represent the mean ± SEM with plots of individual repeats. **p<0.01, ***p<0.001, one-way ANOVA followed by Tukey’s multiple comparisons. (D) A schematic showing signaling pathways activated during TAM polarization. Pathways inhibited by BMS-794833 are shown.

As both BMS-5 and Stattic could inhibit TAM elongation with complementary inhibition of phosphorylated cofilin and STAT3, we tested whether the combination treatment of BMS-5 and Stattic potentiate inhibitor effect on TAM polarization. Combination of BMS-5 and Stattic suppressed TAM elongation more potently than either BMS-5 or Stattic alone at the same dose (5 µM, Figure 6C). The Bliss combination index (Foucquier and Guedj, 2015) was 1.02, suggesting that combination treatment resulted in an additive effect. These results indicate that targeting both LIMK1-actin-related signal and STAT3 better suppress TAM polarization.

Overall, the results above suggest that targeting a single kinase is not sufficient to block TAM polarization. The results reiterate that the potency of BMS-794833 is due to its multi-targeting of SRC, actin regulation, and STAT3, inactivating surrogate pathways that support the plastic nature of TAMs (Figure 6D).

## Discussion

TAMs receive aggregated signals in the form of growth factors, cytokines, chemokines, extracellular vesicles, and metabolites accumulated in TME. How TAMs process this multitude of signals to alter their morphology and function to support the tumor growth and spread remains poorly understood. Current *in vitro* model systems that rely on cytokine-supplemented cultures, such as IL-4 and IL-13, do not adequately recapitulate the complex nature of TME to study TAM function (Wu et al., 2020). To overcome this limitation, we used media collected from tumor tissues, tumor conditioned medium (CM), to induce a complex signaling and phenotypic response through a mixture of secreted factors contained in the CM. Although our *in vitro* system does not model specific cellular interactions that may impact TAM polarization, we confirmed that tumor CM was sufficient to induce pro-tumoral phenotypes known in TAMs. The monocyte cell line and cancer CM-based system provided a relatively uniform and accessible source for TAMs that are compatible with phenotypic drug screening and functional characterization. Using this model system, we could demonstrate interactions between *in vitro*-generated TAMs and (i) cancer cells that led to increased cancer cell growth and motility, (ii) endothelial cells leading to the promotion of microvessel formation, and (iii) T cells causing inhibition of T cell migration. Collectively, we present a physiologically-relevant model system to study TAMs.

To document the signaling cascades that regulate complex TAM function, we performed molecular characterization of our *in vitro*-derived TAMs. We observed alterations of both gene expression and phosphorylation of proteins previously known to be involved in TAM function, including VEGFA, FN1, CXCR4, IL1B(Azizi *et al*., 2018), CCL20(Samaniego et al., 2018), phosphorylated STAT3(Huynh et al., 2019), AKT (Vergadi et al., 2017), and NF-κB(Mancino and Lawrence, 2010). The *in vitro* systems traced the TAM polarization process and allowed us to study time-dependent changes in multiple signaling pathways during the polarization (Figure S4F). Pro-inflammatory transcription factor NF-κB was phosphorylated on day 1, which was then downregulated at later time points, suggesting that tumor CM first induces inflammatory responses in TAMs then shifts towards anti-inflammatory phenotypes. In contrast, STAT3 was unchanged in earlier timepoints but phosphorylated at later timepoints. These results describe serially programmed signaling cascades during TAM polarization and suggest that re-programming this process may offer an opportunity for inhibiting the pro-tumor activity of TAMs.

A combination of rich and uniform sources of *in vitro*-generated TAMs and robust and quantitative cellular elongation phenotype provides a reliable assay for high-throughput screening. To identify compounds that could compromise TAM function, we carried out a chemical inhibitor screen using a set of 85 kinase inhibitors covering most of the kinome. We identified BMS-794833 as a potent suppressor of cellular elongation in multiple cancer models. The mechanism of action of BMS-794833 involves inhibition of phosphorylation status of multiple proteins, including FAK, SRC, p38 MAPK, and STAT3 (Figure 5). BMS-794833 treatment suppresses protumoral functions of TAMs *in vitro* and reduced tumor growth and metastasis *in vivo* (Figure 4B, D), underscoring the potential utility of our model system for phenotypic-based screening in drug discovery.

Several TAM-targeting therapies are currently in clinical studies, including CSF1-CSF1R, either as monotherapy or in combination with conventional treatment to augment therapeutic effects (Kowal et al., 2019; Mantovani et al., 2017). However, clinical studies have shown that monotherapies exhibit moderate effects (Cannarile et al., 2017), indicating the limited efficacy of targeted therapies in a complex environment. Consistently, our data showed that targeting single kinases was ineffective in inhibiting cellular elongation-based TAM polarization (Figure 6A). On the other hand, BMS-794833 exhibits broad polypharmacology, and a combination of LIMK1 inhibitor BMS-5 with STAT3 inhibitor Stattic was able to inhibit TAM polarization efficiently. Collectively, these data suggest simultaneous targeting of multiple signaling pathways in TAM polarization yields more effective treatment options than the use of more selective agents.

Overall, our study highlights the complex interplay of macrophage polarization with components of the TME. Broadly, our study provides the cancer community with a platform that allows analyses of TAMs under near-physiological conditions. As we enter a new era of data-rich cancer biology, one of the primary challenges is integrating the knowledge of cellular and molecular information into a holistic understanding of cancer as a biological system. Quantitative assays and technologies that enable pharmacological assessment with cellular and molecular phenotypes in the physiologically relevant environment, like the one described in this study, will form an essential component of system-based investigations.

## Acknowledgments

This work was supported by grants from the Breast Cancer Research Foundation [BCRF 17-035], the Sidney Kimmel Foundation [Kimmel Scholar Award], and the American Cancer Society (133870-RSG-19-197-01-CDD). NNA is a recipient of the Fred Hutch Interdisciplinary Training Grant in Cancer Research (T32CA080416) and the Japan Society for the Promotion of Science Overseas Research Fellowship. This research was supported by the Comparative Medicine, Scientific Imaging and Flow Cytometry Shared Resources of the Fred Hutch/University of Washington Cancer Consortium (P30 CA015704). We thank Drs. Milka Kostic, Thomas Bello, and Aleena Arakaki, and members of the Gujral lab for helpful comments and suggestions for the manuscript.

## Author Contributions

N.N-A and T.S.G conceived and designed the study. N.A-A performed experiments. N.N-A and T.SG analyzed data and wrote the manuscript. T.S.G. supervised the project, provided resources and funding support.

## Supplementary Information

**Figure S1.**
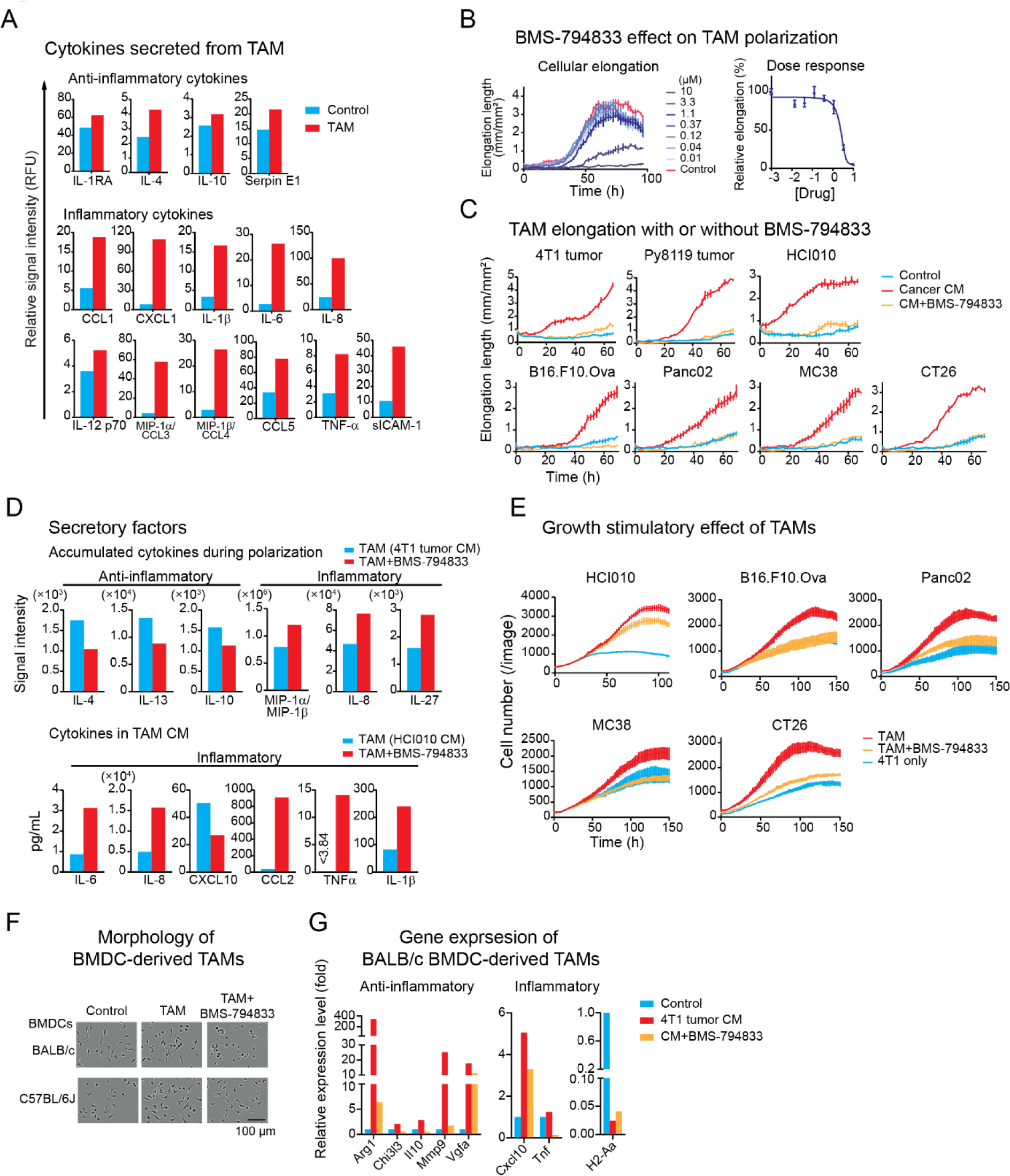
BMS-794833 inhibited TAM polarization with various cancer CM, related to Figures 1, 2, 3. (A) Cytokine profile of in vitro-prepared TAMs. Analyses of cytokines secreted from the THP-1-derived TAMs induced by 4T1 tumor CM. CM collected from the TAM or control macrophages (without tumor CM induction) were analyzed by a cytokine array. The bar graphs show the mean signal intensity from duplicate spots. (B) (Left) Time-course cellular elongation curve and (Right) dose-response curve of TAMs treated with a serial dose of BMS-794833. TAMs were induced with 4T1 cell CM and the dose response at 72 h is shown. The graphs show mean ± SEM of 3 replicates. (C) Time-course TAM elongation induced with CM from breast cancer models (4T1 tumor, Py8119 tumor, PDX (HCI010)) and other cancer types (melanoma, Pancreatic cancer, colon cancer), with or without BMS-794833 at 7.5 µM. (D) Cytokine secretion levels of BMS-794833-treated TAMs. (Left) Accumulated cytokines during THP-1-derived TAM polarization induced with 4T1 tumor CM. Cytokines in the supernatant after 3 days of polarization were measured by a human-specific cytokine array panel. (Right) The absolute concentration of cytokines secreted from TAM CM induced with breast cancer PDX (HCI010) CM. Cytokines in TAM CM were measured by Luminex system. (E) Growth stimulation of 4T1-nucGFP cells by coculturing with TAM polarized with or without BMS-794833 treatment. 4T1-nucGFP were cocultured with BMS-794833-treated TAMs induced by CM from PDX (HCI010), or other cancer models under a reduced serum condition. The growth of 4T1 was evaluated by nuclear GFP counts. The bars indicate mean ± SEM of measurement at each timepoint. Note that the control TAM and 4T1-nuc GFP only data are the same as Figures 2A, B. (F) Representative images of mouse BMDC-derived TAMs treated with BMS-794833. BMDCs from BALB/c or C57BL/6J female mice were induced with M-CSF for 2 days, then cultured with allogenic TNBC tumor CM, 4T1 or Py8119 tumors, supplemented with M-CSF, respectively, for 5 days. BMS-794833 was treated at 7.5 µM together with tumor CM. Images were taken at the endpoint (day 7 from BMDC isolation). (G) TAM marker expressions of BALB/c BMDC-derived TAMs induced with 4T1 tumor CM. Gene expression levels of BMDC-derived TAMs with or without BMS-794833 at 7.5 µM on day 7 were analyzed by qPCR.

**Figure S2.**
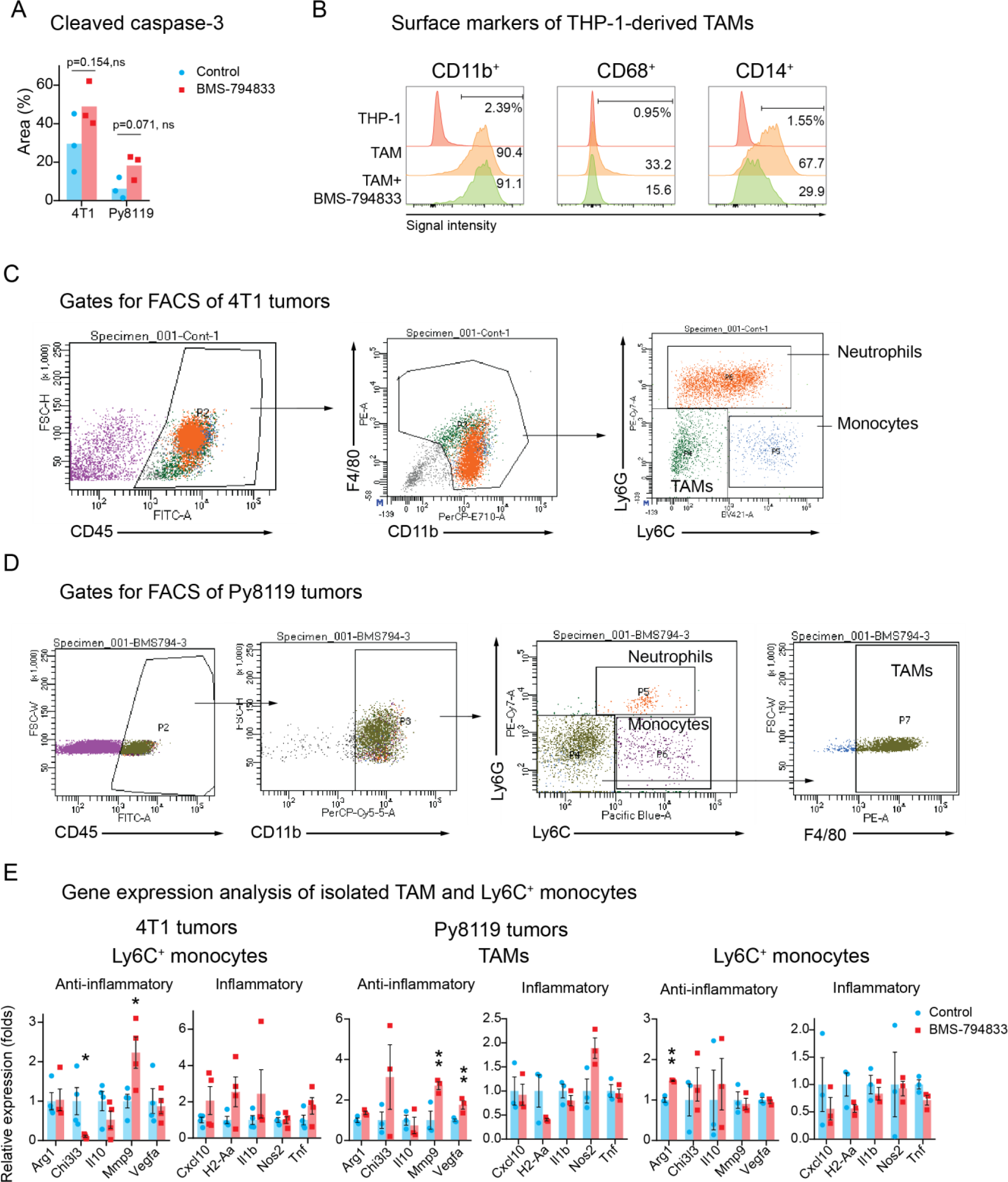
BMS-794833 treatment on mouse TNBC tumors increases apoptosis and modulates characteristics of infiltrated myeloids, related to Figure 4. (A) Immunohistochemistry of cleaved caspase-3 of 4T1 and Py8119 tumors treated with BMS-794833. The representative 3 tumors closest to the average weight at the endpoint were selected from each treatment group of the mouse studies (Figures 4B, C). The percentage of the area positive with cleaved caspase-3 to the entire section was calculated. Statistical analyses were performed with two-tailed unpaired Student’s t-test. (B) Flow cytometry analyses of surface markers of THP-1-derived TAMs. THP-1-derived macrophages were induced with 4T1 tumor CM with or without BMS-794833 at 7.5 µM. All cells were first gated with viability. Histograms of a modal of the total counted cells were shown. (C) Cell sorting gates for isolating TAM, Ly6C^+^ monocytes, and Ly6G^+^ neutrophils from 4T1 tumors. After gated with a viability dye and CD45^+^ cells, CD11b^+^ and/or F4/80^+^ cells were collected, then sorted with Ly6C and Ly6G expression status. (D) Cell sorting gates for isolating TAM, Ly6C^+^ monocytes, and Ly6G^+^ neutrophils from Py8119 tumors. After gated with a viability dye, CD45^+^CD11b^+^ cells were gated, then sorted with Ly6C and Ly6G expression status. TAMs were sorted as F4/80^+^ cells from Ly6C^-^Ly6G^-^ cells. (E) (Left) Gene expression analysis of Ly6C^+^ monocyte population isolated from 4T1 tumors. (Right) Gene expression analysis of TAM and Ly6C^+^ monocyte population sorted from Py8119 tumors. *p<0.05, **p<0.01, unpaired Student’s t-test.

**Figure S3.**
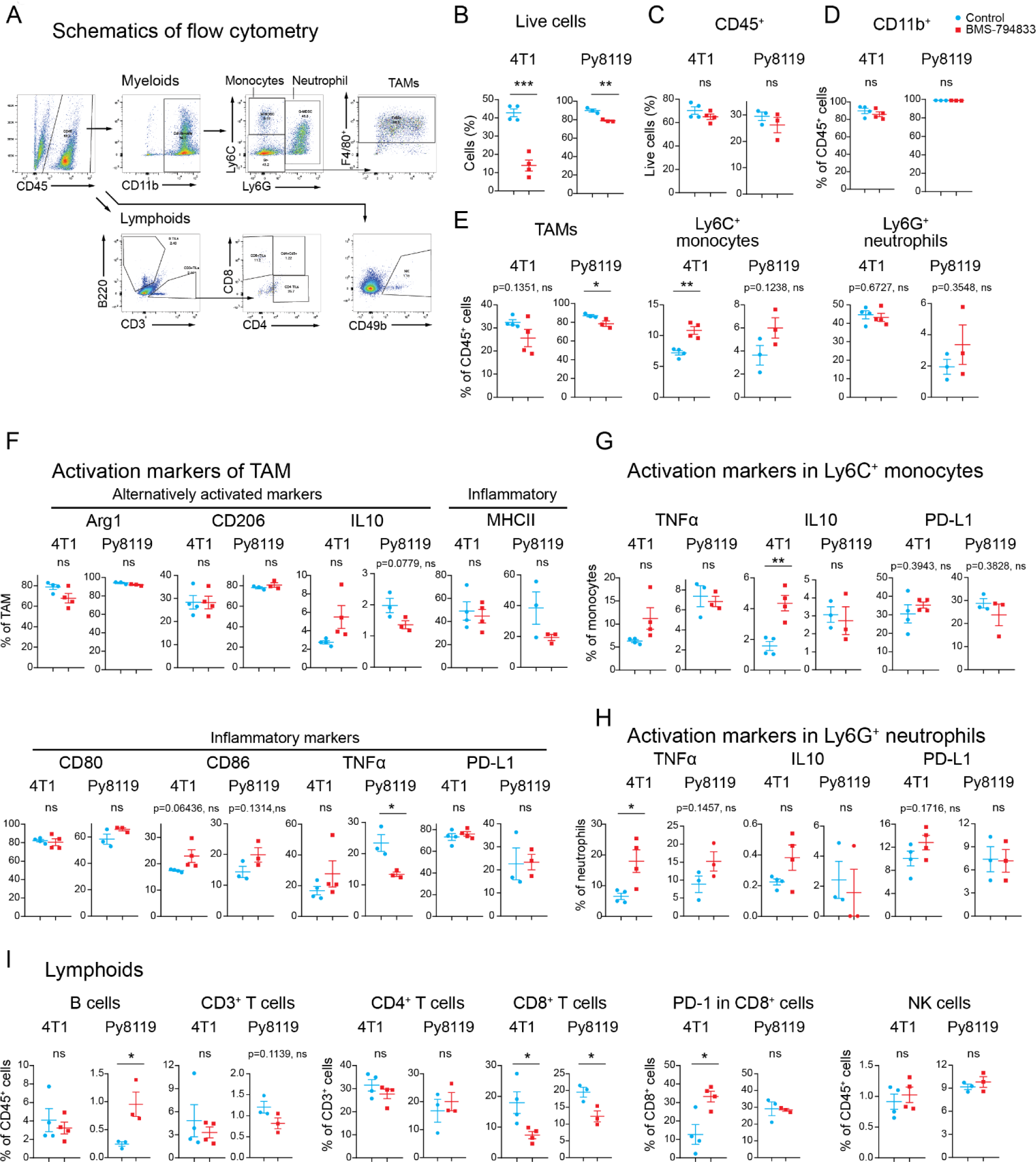
Flow cytometry analyses of immune cell population of TNBC tumors treated with BMS-794833, related to Figure 4. Flow cytometry analyses of immune cell populations of 4T1 mouse tumors (Figure 4B) and of Py8119 mouse tumors (Figure 4C). 4T1 tumors transplanted subcutaneously to BALB/c female mice were treated with BMS-794833 at 25 mg/kg dose biweekly or with vehicle control. Tumors were collected at the experimental endpoint. Tumors from 2 mice were pooled into 1 assay. (A) Gating strategies for flow cytometry analysis. CD45^+^ population was gated after viable cell selection. (B) Live cells. (C) CD45^+^ cells in live cells. (D) CD11b^+^ cells in CD45^+^ cells. (E) TAMs (CD45^+^CD11b^+^Ly6C^-^Ly6G^-^F4/80^+^), monocytes (CD45^+^CD11b^+^Ly6C^+^Ly6G^-^), and neutrophils (CD45^+^CD11b^+^Ly6G^+^) in CD45^+^ cells. (F) TAMs with alternatively activated macrophage and inflammatory macrophage markers. (G) Gene expression of Ly6C^+^ monocytes of the indicated markers. (H) Gene expression of Ly6G^+^ neutrophils of the indicated markers. (I) Lymphoid populations. Error bars indicate mean ± SEM. *p<0.05, **p<0.01, with two-tailed unpaired Student’s t-test.

**Figure S4.**
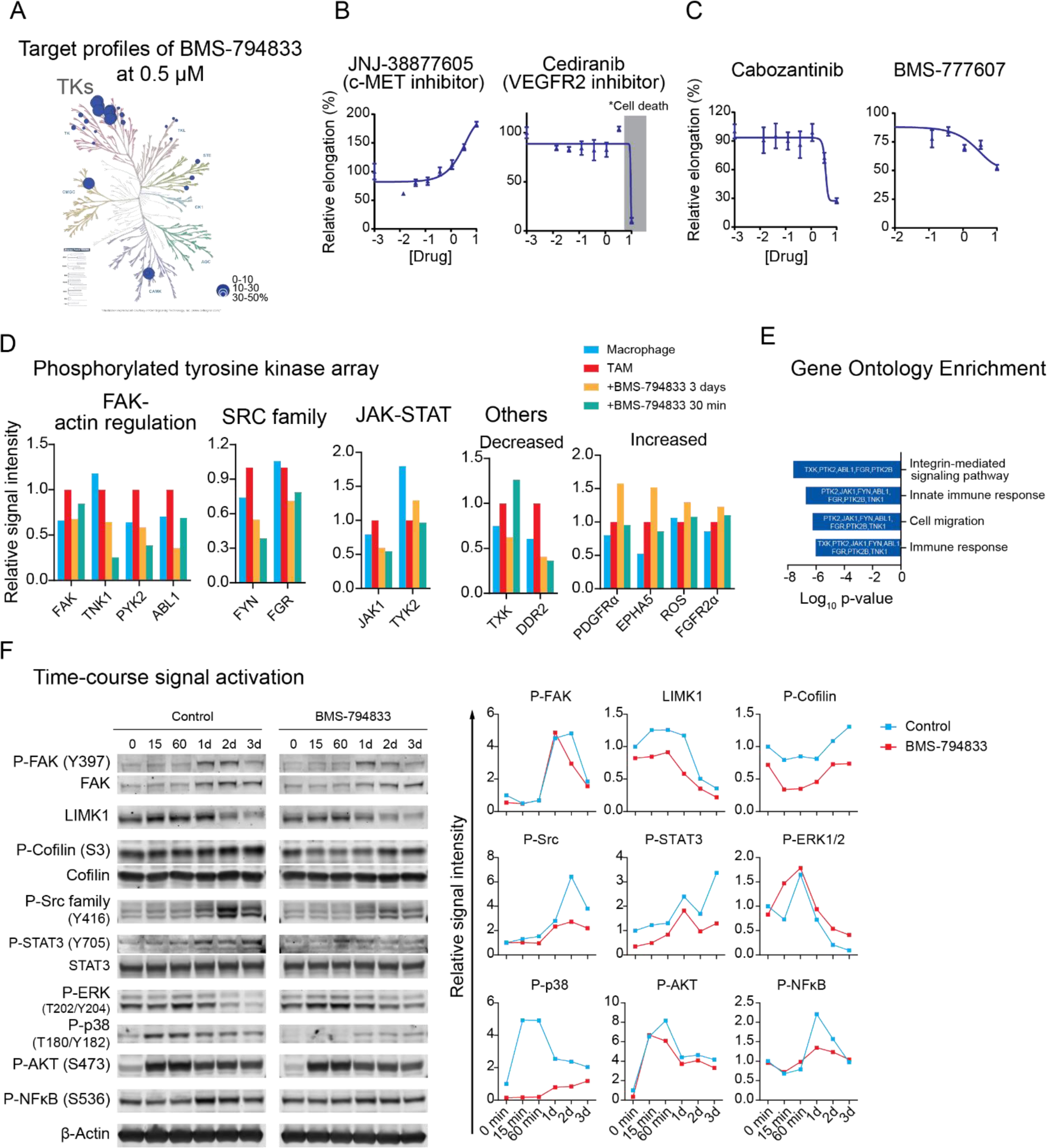
BMS-794833 alters phosphorylation of non-canonical targets during TAM polarization, related to Figure 5. (A) Target kinase profile of BMS-794833 plotted on the kinome tree. The 29 kinases shown in Figure 5A that are inhibited by BMS-794833 at 0.5 µM to below 50% residual activities were plotted. The image was prepared using KinMap (Eid et al., 2017). TKs: tyrosine kinases. (B) Effect of selective inhibitors targeting c-MET and VEGFR2 on cellular elongation of TAMs. (Left) Dose response curve of a selective c-MET inhibitor, JNJ-38877605, which share little structural similarities with BMS-794833. (Right) Dose response curve of a VEGFR2 inhibitor, cediranib. Inhibitors were tested within 0 to 10 µM range and dose response was plotted in log_10_ scale. The concentration causing cellular death is shaded with gray. The plots show the mean ± SEM with n=3. (C) Dose response curves of inhibitors structurally-relevant to BMS-794833, cabozantinib and BMS-777607. The plots show the mean ± SEM with n=3. (D) Altered phosphorylation of kinases in BMS-794833-treated TAM detected by phosphorylated tyrosine kinase panels. THP-1-derived TAM polarized with 4T1 tumor CM were treated with BMS-794833 at 7.5 µM for 3 days or for the last 30 min before sample collection. Macrophages without CM induction were used as control. The average of signals detected from duplicated spots is shown. (E) Gene Ontology enrichment analysis of tyrosine kinases downregulated phosphorylation (below 0.72 folds) after 3 days of BMS-794833 treatment identified in tyrosine kinase array panel. Log_10_ P values of the false discovery rate are shown with enriched genes at each group. (F) Time-course analysis of signal activation during TAM polarization with or without BMS-794833 treatment. (Left) Representative immunoblot images of phosphorylated and total proteins. (Right) Relative quantification of signal intensities. Signal intensities were normalized to control TAM on day 0.

**Figure S5.**
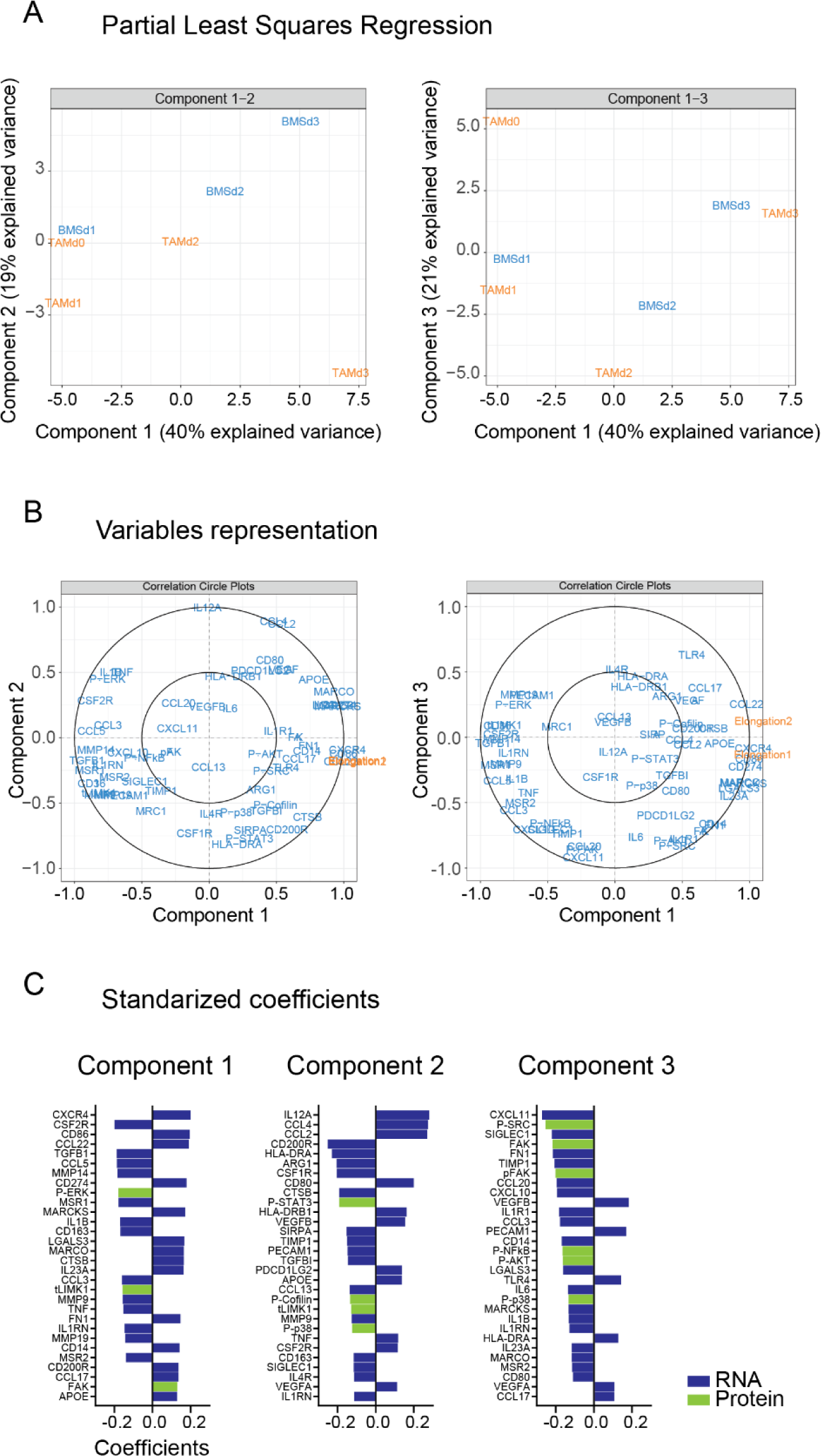
PLS-R analysis segregated factors explaining time-course changes and BMS-794833 treatment during TAM polarization with BMS-794833, related to Figure 3E, Figure S4F. TAMs at different timepoints of the polarization with or without BMS-794833 treatment were analyzed with PLS-R. Time-course gene expression analysis (Figure 3E) and semi-quantified phosphorylated and total protein signal of western blot (Figure S4F) were used as X variates to explain cellular elongation as Y variates. (A) (Left) Biplots of component 1-2 and component 1-3. (B) Distribution of each variable in component 1-2 and component 1-3 space. (C) Top 30 coefficients of variables that contribute to the component 1, 2, and 3 shown in (A). Data from qPCR are colored in blue and the data from WB in green.

**Figure S6.**
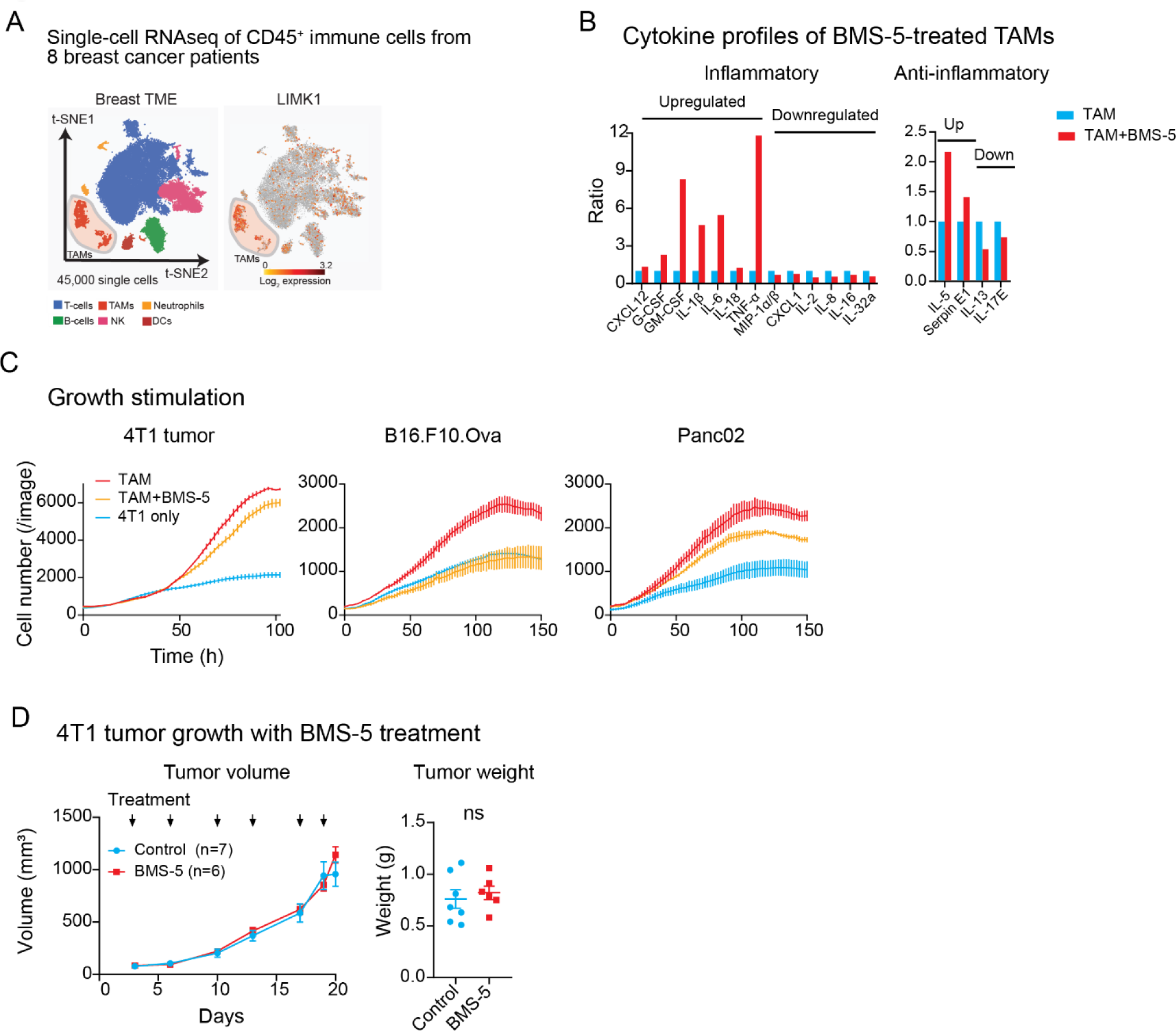
A selective LIMK inhibitor BMS-5 partially suppresses TAM polarization, related to Figure 6. (A) Expression of LIMK1 in population of single-cell RNAseq data of CD45^+^ cells of breast cancer patients. The data from Azizi et al., 2018 were analyzed. (B) Secreted cytokines from BMS-5-treated TAMs. Accumulated cytokines secreted during the polarization of BMS-5-treated TAMs with 4T1 tumor CM for 3 days were analyzed. A human-specific cytokine array panel was used to detect only cytokines only secreted from THP-1-derived TAMs. (C) Growth stimulatory effect on 4T1 breast cancer cells by coculturing with TAM treated with BMS-5 at 10 µM. 4T1-nucGFP cells were cocultured with TAMs induced with CM from 4T1 tumor, melanoma (B16.F10.Ova), or pancreatic cancer (Panc02) under reduced serum condition. The growth of 4T1 was evaluated by nuclear GFP counts. Note that the control TAM and 4T1 only data are the same as Figure 3F and 2B. (D) 4T1 tumor growth treated with BMS-5. BALB/c female mice bearing subcutaneous 4T1 tumor were treated biweekly with BMS-5 at 30 kg/mg or vehicle control with intratumoral injections. (Left) time-dependent tumor volume. The timing of drug administration is shown in black arrows. (Right) Tumor weight at the experimental endpoint. Statistical analyses were performed with two-tailed unpaired Student’s t-test.

## Description of Supplementary Tables

***Table S1 List of 85 kinase inhibitors used for screening on THP-1-derived TAM and their response, related to Figure 3, STAR methods***

Column C denotes the maximum concentration of inhibitor that causes a change in elongation without affecting cell viability. Column D indicates the maximum altered cellular elongation levels observed compared to control TAMs shown as a percentage (control = 0%). Negative values mean decreased elongation, positive values mean increased elongation. The inhibitors that caused cellular deaths were evaluated as no change in elongation or omitted from the analysis. The results are summarized in Figure 3C.

***Table S2 Target kinase profile of BMS-794833 at 0.5 µM, related to Figure 5, Figure S4A***

The residual activities of 298 kinases under the presence of BMS-794833 at 0.5 µM in acellular system were shown in the percentage. Control is 100 %. The data was extracted from Rata, et al, 2020.

***Table S3 List of antibodies used for flow cytometry analyses, related to STAR methods***

The antibodies were used in flow cytometry and cell sorter analysis of Figure 4E, Figs. S5, S6, S7.

